# The mitochondrial-targeted antioxidant SkQ1 prevents mitochondrial-linked apoptosis but not necroptosis or skeletal muscle atrophy in ovarian cancer

**DOI:** 10.1101/2024.10.22.617245

**Authors:** Shahrzad Khajehzadehshoushtar, Luca J. Delfinis, Fasih A. Rahman, Madison C. Garibotti, Shivam Gandhi, Aditya N. Brahmbhatt, Brooke A. Morris, Bianca Garlisi, Sylvia Lauks, Caroline Aitken, Zhu-Xu Zhang, Jeremy A. Simpson, Joe Quadrilatero, Jim Petrik, Christopher G.R. Perry

**Author notes:** Email Addresses,. <u>Address for Correspondence:</u> Christopher Perry, PhD, School of Kinesiology and Health Science, Muscle Health Research Centre, 344 Norman Bethune College, York University, 4700 Keele Street, Toronto, Ontario M3J 1P3 (P).

## Abstract

The degree to which mitochondrial-linked cell death pathways contribute to skeletal muscle atrophy during cancer remain unknown. Here, we combined a novel and robust mouse model of metastatic ovarian cancer with chronic administration of the mitochondrial-targeted antioxidant SkQ1 to determine the time-dependent and muscle-specific relationships of mitochondrial-linked apoptosis and necroptosis to the development of muscle atrophy in the type II B-rich gastrocnemius. Early-stage ovarian cancer reduced type II B fibre cross-sectional area in the gastrocnemius but did not alter mitochondrial H_2_O_2_ emission despite increased activities of mitochondrial-linked caspase-9 and-3 regulators of apoptosis. During late-stage ovarian cancer, sustained atrophy was associated with increased mitochondrial H_2_O_2_ emission potential *in vitro*, a greater probability of calcium-triggered mitochondrial permeability transition and increases in downstream caspase-9 and −3 activity. SkQ1 attenuated mitochondrial H_2_O_2_ emission and caspase-9 and −3 activity in late-stage ovarian cancer but did not prevent atrophy. Necroptosis markers were heterogeneous across time with total RIPK1 increasing during early-stage cancer which reverted to normal levels by late-stages while phosphorylated RIPK3 decreased below control levels. These discoveries indicate that preventing increases in mitochondrial-linked apoptotic caspase-9 and −3 activity during late-stage ovarian cancer with SkQ1 does not prevent atrophy of type II B fibres. Furthermore, necroptotic markers are inconclusive during cancer in this muscle type but are not modified by SkQ1. These results do not support a causal relationship between mitochondrial H_2_O_2_-linked apoptosis or necroptosis and atrophy in type II B fibres during ovarian cancer but do not rule out potential relationships in other muscle types.

**Key Points:**

1. Cancer increases mitochondrial reactive oxygen species (ROS) in skeletal muscle during atrophy, but the role of ROS in regulating cell death remains unknown.
2. We show that attenuating gastrocnemius mitochondrial ROS with the mitochondrial-targeted antioxidant SkQ1 prevented mitochondrial-linked pro-apoptotic caspase 9- and 3-activities but did not affect markers of necroptosis in a mouse model of ovarian cancer.
3. Reductions in gastrocnemius muscle fibre cross-sectional areas and the wet weights of several muscles were not prevented by SkQ1.
4. These findings demonstrate that mitochondrial ROS regulate apoptotic caspases but not necroptosis, and neither pathway is linked to gastrocnemius atrophy in mice with ovarian cancer.
5. The degree to which mitochondrial ROS-linked cell death pathways regulate muscle mass in other muscle types and cancer models requires further investigation.

## Introduction

Muscle mass is a significant predictor of cancer survival (Martin *et al*., 2013). Approximately 70% of cancer patients develop cancer cachexia, a syndrome characterized by an irreversible loss of muscle mass (atrophy) and often adipose tissue, that is irreversible with nutritional intervention (Fearon *et al*., 2011). Skeletal muscle atrophy in cancer patients correlates with greater dose-limiting toxicity, increased likelihood of treatment discontinuation, and prolonged hospital stays (Prado *et al*., 2009; Van Vledder *et al*., 2012; Poisson *et al*., 2021). Furthermore, skeletal muscle loss during cancer is linked to lower survival rates following tumour resection surgeries, a higher likelihood of tumour recurrence, and diminished overall survival rates, contributing to nearly 30% of cancer-related fatalities globally (Fearon *et al*., 2011). Despite its prevalence and impact on individuals’ lives, there is no clinically available therapy that directly prevents or treats cachexia. Thus, a comprehensive understanding of the molecular mechanisms driving muscle atrophy during cancer progression is essential for developing effective therapeutic strategies.

Muscle atrophy during cancer cachexia is thought to be driven by circulating factors produced during cancer that promote protein degradation and myofibrillar protein loss through mechanisms like the ubiquitin-proteasome system (UPS) and autophagy (Martin *et al*., 2023). However, emerging research indicates that skeletal muscle mitochondrial stress develops during cancer cachexia and may directly contribute to muscle atrophy (Gilliam *et al*., 2016; Brown *et al*., 2017, 2020; Smuder *et al*., 2020; Pin *et al*., 2022; Delfinis *et al*., 2022, 2024). In particular, excessive mitochondrial reactive oxygen species in the form of hydrogen peroxide emission (mH_2_O_2_) in the plantaris, diaphragm, quadriceps, and tibialis anterior (TA) muscles of various cancer models is seen before or concurrent with atrophy (Gilliam *et al*., 2016; Brown *et al*., 2017; Delfinis *et al*., 2022, 2024). Additionally, pharmacological prevention of excessive mH_2_O_2_ using the cardiolipin-targeting peptide SS-31 has shown promise in mitigating muscle mass loss and improving overall body weight in the C26 colorectal cancer mouse model (Smuder *et al*., 2020; Ballarò *et al*., 2021). Recently, our group demonstrated that SkQ1 partially prevented atrophy in male mice with C26 cancer whereas it caused greater atrophy in female mice (Tsitkanou *et al*., 2024). Likewise, the mitochondrial-targeted antioxidant and quinone analogue MitoQ prevented atrophy in a cell culture model of cancer cachexia (Pin *et al*., 2022). However, mice expressing the H_2_O_2_ scavenging antioxidant catalase within mitochondria were not protected from breast cancer-induced loss of mass in the soleus (Gilliam *et al*., 2016). Thus collectively, the mechanisms by which mitochondrial-targeted compounds prevent atrophy remains unknown.

The mechanisms by which mitochondrial ROS contribute to skeletal muscle atrophy during cancer are complex. With growing evidence of the involvement of mitochondrial ROS in regulated cell death pathways, it is plausible that these pathways mediate the link between mitochondrial ROS and muscle atrophy. Two prominent pathways in this context include apoptosis and necroptosis. In the former, excess mitochondrial ROS and calcium trigger the formation of the mitochondrial transition pore (mPTP) whereby increased permeability of the double membrane system causes release of pro-apoptotic factors through a ‘mitochondrial permeability transition’ (mPT) event (Bonora *et al*., 2021; Bernardi *et al*., 2023). These factors activate caspase-9 and −3, myofibrillar protein degradation, and cell death (Du *et al*., 2004; Adhihetty *et al*., 2007; Skinner *et al*., 2021). Caspase-9 and −3 as well as DNA fragmentation are also elevated in muscle biopsies from patients with gastric, colorectal, and gastrointestinal cancers and rodent models but the role of mPT in this process is unknown (Belizário *et al*., 2001; Busquets *et al*., 2007; Baltgalvis *et al*., 2010; de Castro *et al*., 2019). Necroptosis has been explored in multiple cell types as well as muscle disorders such as polymyositis and Duchenne muscular dystrophy but to our knowledge, has never been assessed during cancer (Morgan *et al*., 2018; Gan *et al*., 2019; Oikonomou *et al*., 2021; Kamiya *et al*., 2022). In this form of cell death, circulating cytokines like Tumour necrosis factor (TNF) trigger the autophosphorylation of Receptor interacting serine/threonine kinase 1 (RIPK1), which binds and phosphorylates RIPK3 to form the ‘necrosome’ (Huang *et al*., 2013; Gan *et al*., 2019; Khoury *et al*., 2020; Chu *et al*., 2021; Oikonomou *et al*., 2021). The necrosome phosphorylates mixed-lineage kinase domain-like (MLKL) pseudokinase monomers, leading to pore formation on the cell membrane and subsequent cell death (Rodriguez *et al*., 2016; Samson *et al*., 2020). mH_2_O_2_ also promotes necroptosis by oxidizing RIPK1, further facilitating necrosome-mediated membrane permeabilization (Zhang *et al*., 2017). However, a direct relationship between mitochondrial ROS and both apoptosis and necroptosis has not been investigated in cancer cachexia.

It has been proposed that pre-clinical development of therapies to treat cancer cachexia would have greater translational potential to humans if assessments were performed in animal models whereby cancer is induced in the host organ during adulthood in immunocompetent mice that demonstrate metastasis (Mueller *et al*., 2016; Penna *et al*., 2016; Tomasin *et al*., 2019). In this regard, we recently reported a novel and robust model of ovarian cancer that is induced during adulthood in immunocompetent mice through injection of syngeneic cancer cells in the ovarian bursa (Delfinis *et al*., 2024). This model resulted in a progressive development of ovarian cancer over approximately three months including ascites fluid (a hallmark of stage III cancer) and metastasis observed in the abdominal cavity at 75 and 90 days, respectively. Reduced limb muscle fibre cross-sectional areas were observed at 75 days while lower overall muscle mass was observed at 90 days with cancer. Reduced skeletal muscle fibre cross-sectional area at 75-days accompanied increases in mH_2_O_2_. These findings support the use of this model for continued exploration of mitochondrial contributions to cancer as well as exploration of new therapeutics.

The objective of this study was to investigate the role of mH_2_O_2_-mediated regulated cell death pathways, apoptosis and necroptosis, in cancer-induced muscle atrophy. To achieve this objective, we examined whether attenuating mH_2_O_2_ through a commercially-available mitochondrial-targeting antioxidant (SkQ1) could prevent regulated cell death and preserve muscle mass in cancerous mice. While the theoretical link between cell death and atrophy seems plausible, we did not form a hypothesis given the lack of prior knowledge regarding these cell death pathways in cancer cachexia models (Glass, 2010). Using an orthotopic metastatic epithelial ovarian cancer (EOC) model of stage III ovarian cancer (Delfinis *et al*., 2024), we chose to assess the white gastrocnemius muscle given i) we previously demonstrated a loss in muscle mass in this model, ii) it is rich in type IIB fibres which has shown a greater reduction in cross sectional area compared to other fibre types across different muscles in the EOC model, and iii) it provides sufficient tissue mass to perform the majority of our experimental measures within the same muscle type. We found that atrophy surprisingly preceded elevated mH_2_O_2_ in the white gastrocnemius muscle, and attenuating mH_2_O_2_ with SkQ1 did not prevent atrophy. Our assessment of apoptotic and necroptotic markers revealed heterogeneous and divergent responses, with apoptosis markers increasing and necroptotic markers remaining unchanged relative to non-cancer mice during EOC progression. Furthermore, SkQ1 lowered caspase-9 and −3 and certain necroptotic markers despite having no effect on atrophy, highlighting that mitochondrial-linked apoptosis and necroptosis do not contribute to atrophy in type II B-rich muscle during ovarian cancer.

## Methods

### Study Design

Transformed murine ovarian surface epithelial cells (EOC, ID8 cells) from C57BL/6J mice were injected beneath the ovarian bursa to establish an orthotopic, syngeneic mouse model of EOC (Greenaway *et al*., 2008, 2009; Russell *et al*., 2014; Matuszewska *et al*., 2019; Delfinis *et al*., 2024). These were compared to phosphate-buffered saline (PBS) injected controls (Control, n=24). Mice in all groups received initial PBS or EOC ID8 cells at slightly staggered ages in order to ensure a sustainable rate of endpoint assessments given the requirement for fresh tissue for a number of experimental assays listed below. EOC tumour-bearing mice were assessed at early-stage (35-46 days) and late-stage (58-84 days) post tumour injections. Control mice were assessed 91-105 days post PBS injections. Time point ranges were partially informed by our previous work that reported similar ages for pre-atrophy weakness and atrophy in this mouse model (Delfinis *et al*., 2024). Immediately following EOC injections and for the duration of the study, half of the mice received 250 nmol/kg body weight SkQ1 in their drinking water (Early-EOC-SkQ1, n=12; Late-EOC-SkQ1, n=30), while the rest received standard water (Early-EOC, n=12; Late-EOC, n=30). A higher sample size was required for Late-EOC mice given the unpredictability, heterogeneity, and complications of cancer at this time. Mice in the Late-EOC groups were aged to the latest timepoint possible prior to presentation of combinations of the following criteria: 10% body weight loss, > 20mL of ascitic volume collected during paracentesis, > 5 ascites paracentesis procedures, and/or subjective changes in behavioural patterns consistent with removal criteria as per animal care guidelines (self-isolation, ruffled fur/poor self-grooming and irregular gait) similar to our previous research. During the duration of the study, 6 Late-EOC and 4 Late-EOC-SkQ1 mice expired naturally and were included in our survival analysis listed below.

### Animal Care and Tumour Injection

#### Cell Culture

Spontaneously transformed murine ovarian epithelial cells (ID8), donated by Drs. K. Roby and P. Terranova from Kansas State University, were cultured in Dulbecco’s Modified Eagle Medium (DMEM) supplemented with 10% Fetal Bovine Serum (FBS) and 1% antibiotic/antimycotic (Greenaway *et al*., 2008, 2009; Russell *et al*., 2014; Matuszewska *et al*., 2019; Delfinis *et al*., 2024).

#### Animals and Tumour Implantation

Female C57BL/6 mice, aged 10 and 16 weeks, were injected with ID8 cells at the University of Guelph in accordance with the Canadian Council on Animal Care following the methodology previously outlined previously (Delfinis et al., 2024; Ogilvie et al., 2024). Under isoflurane anesthesia, a dorsal incision was made, and 1.0 x 10^6^ ID8 cells suspended in 5 µl sterile PBS were injected directly beneath the bursa of the left ovary. Control mice underwent an identical procedure but received a 5 µl injection of sterile PBS without cancer cells. The mice were observed for 2 weeks before being transferred to the animal facility at York University, where they were housed for the remainder of the study in accordance with the Canadian Council on Animal Care.

Mice were housed in groups of four, and weekly measurements of food and water intake were recorded until the endpoint. Body weights were initially measured weekly for 45 days, then daily due to the rapid deterioration in the animals’ condition. Mice displaying 10% body weight loss, > 20mL of ascitic volume collected during paracentesis, > 5 ascites paracentesis procedures, and/or subjective changes in behavioural patterns consistent with removal criteria as per animal care guidelines (self-isolation, ruffled fur/poor self-grooming and irregular gait) were euthanized promptly.

#### SkQ1 Administration

Immediately following tumour implantation, mice in the SkQ1 groups received SkQ1 (Cayman Chemical, 19891, dissolved in a 1:1 ethanol and water solution) in their drinking water (250 nmol/kg body weight/day) until the endpoint (Anisimov *et al*., 2011; Fedorov *et al*., 2022). Also, while we did not compare dosages of SkQ1 in this study, the dosage we chose was similar to other studies that showed positive benefits in other disease models (Anisimov *et al*., 2011; Fedorov *et al*., 2022; Tsitkanou *et al*., 2024). Mice were estimated to weigh 25g on average and drink 5 mL/day, resulting in a 1.25 nmol/mL concentration of SkQ1. The ethanol concentration in the 1.25 nmol/mL drinking water was approximately 0.0008%, derived from a 50 mg/mL 1:1 ethanol stock. Thus, non-SkQ1 groups (Early- and Late-EOC) received regular drinking water, as the ethanol concentration was negligible (0.008%).

#### Abdominal Paracentesis

Primary tumours emerged approximately 8.5 weeks post-injection of EOC ID8 cells, followed by ascites development, resembling stage III EOC. To alleviate discomfort and ensure proper functioning of nearby organs, mice with significant ascites underwent abdominal paracentesis using a 25-gauge needle under isoflurane anesthesia.

#### Surgical Procedures

Mice were euthanized under isoflurane and hearts were removed. Various tissues, including the quadriceps (Quad), gastrocnemius (Gastroc), tibialis anterior (TA), extensor digitorum longus (EDL), plantaris, soleus, heart, spleen, kidneys, liver, lungs, tumour, and inguinal fat, were collected. Muscle tissues, spleen, tumour, and inguinal fat were weighed, and all tissues were snap-frozen in liquid nitrogen and stored at −80°C. The red portions of the gastrocnemius muscles were excised and similarly snap-frozen. For half of the animals, the white gastrocnemius (WG) was divided: one piece was used for bioenergetic assays, while the other was used for sub-cellular fractionation (not used for this study). For the second half of mice per group, one WG was flash-frozen, and the other was embedded in optimal cutting temperature (OCT) medium and frozen. Due to the lower n size of the early-stage cancer groups, half of one gastrocnemius was used for bioenergetics (only white gastrocnemius), while the other half was OCT-embedded (mixed gastrocnemius). The second gastrocnemius (only white portion) was used for western blotting. A similar approach was used for the quadriceps muscle, with two quadriceps utilized for sub-cellular fractionation, one snap-frozen, and the other embedded in OCT.

### Mitochondrial Bioenergetic Assessments

#### Preparation of permeabilized muscle fibres

Preparation of permeabilized muscle fibre bundles (PmFB) and H_2_O_2_ analysis was performed as previously established in our lab (Hughes *et al*., 2019; Delfinis *et al*., 2022; Bellissimo *et al*., 2023). Briefly, WG was immediately placed in BIOPS comprising (mM) 50 MES Hydrate, 7.23 K_2_EGTA, 2.77 CaK_2_EGTA, 20 imidazole, 0.5 dithiothreitol, 20 taurine, 5.77 Na_2_ATP, 15 Na_2_PCr, and 6.56 MgCl_2_.6 H_2_O (pH 7.1) following removal. Under 10x magnification, WG was trimmed of fat and connective tissue, and divided into 9 smaller bundles (8 for H_2_O_2_ analysis and 1 for calcium retention capacity). All bundles were made from the centre of the white gastrocnemius muscle and gently separated along the longitudinal axis.

Bundles were then treated with 40 μg/ml saponin in BIOPS on a rotor for 30 minutes at 4 °C (permeabilization). For bundles intended for mH_2_O_2_ measurement related to Complex I, II, and pyruvate dehydrogenase complex (PDC), an additional treatment with 35 μM 2,4-dinitrochlorobenzene (CDNB) was applied during the permeabilization step. This step served to deplete glutathione, enabling detectable rates of mH_2_O_2_.

Following permeabilization, bundles were divided into 2 groups: 1) bundles intended for mitochondrial H_2_O_2_ emission (mH_2_O_2_) were placed in buffer Z comprising (in mM) 105 K-MES, 30 KCl, 10 KH_2_PO_4_, 5 MgCl_2_ · 6H_2_O, 1 EGTA and 5 mg/mL BSA (pH 7.1) on a rotator at 4 °C for 15 minutes. 2) bundles intended for calcium retention capacity were placed in 1 mM EGTA in Buffer Y comprising (mM) 250 sucrose, 10 Tris-HCl, 20 Tris-base, 10 KH_2_PO_4_, 0.5 mg/ml BSA (pH 7.2) on a rotator for 10 minutes at 4 °C. Subsequently, CRC bundles were transferred to a second wash of Buffer Y with 10 μM Blebbistatin (BLEB, to prevent ATP induced muscle contraction) on a rotor at 4 °C for a minimum of 10 minutes until measurements were initiated.

#### Mitochondrial H_2_O_2_ Emission (mH_2_O_2_)

mH_2_O_2_ was determined fluorometrically (QuantaMaster 40, HORIBA Scientific, Edison, NJ, USA) in a quartz cuvette with continuous stirring at 37 °C, in 1 ml Buffer Z supplemented with 10 μM Amplex Ultra Red, 0.5 U/mL horseradish peroxidase, 1 mM EGTA, 40 U/mL Cu/Zn-SOD1, 5 μM BLEB, and 20 mM Cr. Site specific induction of H_2_O_2_ was measured through the addition of either 10 mM Pyruvate and 2 mM malate (NADH, complex I), 10 mM succinate (FADH_2_, complex I via reverse electron flux from complex II) or 2.5 μM Antimycin A (Complex III). Additionally, using 0.5 μM rotenone, a complex I inhibitor, with 10 mM pyruvate, electron slip specific to pyruvate dehydrogenase complex was also measured in CDNB-treated fibres. Following the induction of maximal mH_2_O_2_ by complex I and II substrates, ADP was titrated progressively to attenuate mH_2_O_2_ and model kinetics during oxidative phosphorylation. The rate of mH_2_O_2_ was calculated from the slope (F/min) of a standard curve established with the same reaction conditions and normalized to fibre bundle dry weight.

#### Mitochondrial Calcium Retention Capacity (CRC)

CRC was determined fluorometrically (QuantaMaster 80, HORIBA Scientific, Edison, NJ, USA) in a quartz cuvette with continuous stirring at 37 °C. Prior to the initiation of each experiment, the cuvette was placed on a stir plate with 500 μl water and 10 mM EGTA. The water was then aspirated from the cuvette but not rinsed, leaving the EGTA coating on the cuvette walls to chelate any residual Ca^2+^ in the assay buffer. Background fluorescence was obtained following the addition of 5 mM glutamate, 2mM malate, 25 μM ATP and the PmFB to 300 μl CRC buffer comprising 1 μM Calcium Green-5N (Invitrogen), 20 mM creatine, 40 μM EGTA, 5 μM BLEB, 2 μM thapsigargin in Buffer Y. Calcium uptake was then initiated by a single 8 nmol pulse of CaCl_2._ Subsequent 4 nmol pulses of CaCl_2_ were added until mitochondrial permeability transition pore (mPTP) opening was evident. Three 0.5 mM pulses of CaCl_2_ were then added to saturate the fluorophore and establish a fluorescence maximum (Fmax). Changes to free Ca^2+^ in the cuvette during mitochondrial Ca^2+^ uptake were calculated using the known K_d_ for Calcium Green-5N and the equations established for calculating free ion concentrations using ion sensitive fluorophores.

Upon completion of mH_2_O_2_ and CRC experiments, the fibres were lyophilized in a freeze-dryer (Labconco, Kansas City, MO, USA) for > 4 hours and weighed on a microbalance (Sartorius Cubis Microbalance, Gottingen, Germany).

### Validating Necroptosis Antibodies

It was crucial to verify the reliability of the purchased necroptosis antibodies in muscle tissue due to the scarcity of literature on this cell death mechanism in skeletal muscle. Initially, we induced necroptosis in C2C12 myoblasts, kindly donated by Dr. Olasunkanme Adegoke from York University, using a well-established cell culture necroptosis model (Gan *et al*., 2019; Kim *et al*., 2021). Additionally, RIPK3 knockout (KO, Genentech, Inc., South Sanfrancisco, CA, USA) gastrocnemius and quadriceps were used as negative controls (Newton *et al*., 2004). We also used gastrocnemius and quadriceps muscles from our D2.*mdx* model of Duchenne Muscular Dystrophy (DMD) tissue bank as an additional positive control, as recent literature has demonstrated necroptosis D2.*mdx* mice (Morgan *et al*., 2018; Bellissimo *et al*., 2023).

#### C2C12 treatment

C2C12 cells were maintained in growth medium (GM) consisting of DMEM, 10% FBS and 1% penicillin/streptomycin. C2C12 myoblasts were treated with TNF to activate the cell death mechanisms. A pan-caspase inhibitor, z-VAD-FMK, was used to inhibit apoptosis. Additionally, a caspase-8 specific inhibitor z-IETD-FMK was used as a secondary measure to further inhibit apoptosis. Finally, to induce necroptosis, BV6, a general inhibitor of proteins that ubiquitinate and reduce RIPK1’s pro-necroptotic activity was employed to promote RIPK1-mediated necroptosis. Briefly, C2C12 myoblasts, seeded on 6-well culture plates, were exposed to a combination of 10 nM/ml mouse TNF (T, Peprotech #315-01A), 1 μM BV6 (B, APEXBIO #B4653), 20 μM z-Vad-FMK (Z, APEXBIO #A1902), and 20 μM z-IETD-FMK (I, APEXBIO #B3232-1). C2C12 cells were assessed at various time points following TBZI treatment, and at both low-dose (10 nM/ml) and high-dose (100 nM/ml) TNF-α concentrations to optimize the necroptosis model (Kearney *et al*., 2015; Kim *et al*., 2021). It was determined that 10 nM/ml TNF-α, assessed 24 hours following TBZI treatment, yielded the greatest positive results for necroptosis proteins, as evidenced by Western blot analysis. After the treatment period, cells were harvested, homogenized, and subjected to Western blot analysis to confirm the presence and reliability of necroptosis-related proteins.

### Western Blotting

An aliquot of frozen Quadriceps and WG were homogenized in ice cold homogenization buffer comprising (in mM): 10 Tris-HCl, 150 NaCl, 1 EDTA, 1 EGTA, 2.5 Na_4_O_7_P_2_, and 1 Na_3_VO_4_, 1% Triton X-100, 1:200 protease inhibitors (MilliporeSigma #P8340, pH 7.0), and phosphatase inhibitors (MilliporeSigma, #4906845001) using a polytron homogenizer at low speed.

Protein concentrations of homogenized tissues were determined using Bicinchoninic acid assay (BCA, Life Technologies, Carlsbad, CA, USA). 15-60 μg of denatured and reduced protein was subjected to 8-12% gradient SDS-PAGE followed by transfer to low-fluorescence polyvinylidene difluoride membranes (LF PVDF, BIORAD). Membranes were blocked with LICOR Odyssey TBS Blocking Buffer (LI-COR) and immunoblotted 24-48 hours (4°C) with antibodies specific for each protein. The following mouse primary antibodies were used to detect necroptosis: RIPK1 and phospho-RIPK1 (Cell Signalling #3493 and #53286, 79 kDa, 1:500 and 1:250), RIPK3 (ProSci Inc. #2283, 53 kDa, 1:500), MLKL (Abcepta #AP14272, 54 kDa, 1:500), phospo-RIPK3 and phospho-MLKL (Abcam #EPR9516(N)-25 and #EPR9515(2), 53 and 54 kDa, 1:250 and 1:250). A rodent OXPHOS antibody cocktail was used to detect electron transport chain protein subunits (Abcam #ab110413, 1:250): ATP5A (complex V, 55 kDa), UQCRC2 (Complex III, 48 kDa), MTCO1 (Complex IV, 40 kDa), SDHB (Complex II, 30 kDa) and NDUFB8 (Complex I, 20 kDa).

After incubation in primary antibodies, membranes were subjected to three 5-minute washes in TBS-Tween followed by a 1-hour incubation at room temperature with an infrared fluorescent secondary antibody (LI-COR, IRDYE 680RD goat-anti-mouse #925-68070 and goat-anti-rabbit #926-68071) at a previously optimized dilution (1:20,000). Immunoreactive proteins were detected using infrared imaging (LI-COR CLx; LICOR) and subsequently quantified via densitometry using ImageJ (NIH). All images were normalized to Amido Black total protein stain (Sigma, A8181).

### Caspase Activity

Caspase activity assays were conducted as outlined previously (Rahman *et al*., 2024). Briefly, WG muscles were homogenized as above in the absence of protease inhibitors. Samples were incubated at room temperature for 2 hours with 20μM Ac-DEVD-AFC (Alexis Biochemicals) for caspase-3 or Ac-LEHD-AFC (Alexis Biochemicals) for caspase-9 in assay buffer (20 mM HEPES pH 7.4, 10 mM DTT, and 10% glycerol). Fluorescence was measured at room temperature using a Cytation 5 Imaging Multi-Mode Reader (BioTek) with excitation and emission wavelengths at 400 and 505 nm, respectively. The activity assay was normalized to protein concentration of samples (BCA assay) and expressed as fold changes in fluorescence.

### Immunofluorescence

#### Sample Preparation

Quadriceps and full gastrocnemius muscles embedded in O.C.T medium (Thermo Fisher Scientific) were cut into 10 μm sections with a cryostat (HM525 NX, Thermo Fisher Scientific) at −20 °C. Cryosections were used for immunofluorescence analysis.

#### Immunofluorescence Analysis

Muscle fiber type analysis followed the methodology described previously, allowing determination of fiber-type-specific atrophy (Delfinis *et al*., 2022, 2024). Briefly, slides were blocked with 5% goat serum (MilliporeSigma) in PBS for 1 hour at room temperature. Next, slides were incubated with primary antibodies (Developmental Studies Hybridoma Bank, University of Iowa) against MHC I (BA-F8; 1:25), MHC IIa (SC-71; 1:1,000), and MHC IIb (BF-F3; 1:50) for 2 hours at room temperature. Slides underwent three 5-minute PBS washes and incubated with secondary antibodies (Invitrogen, Thermo Fisher Scientific): MHC I; Alexa Fluor 350 IgG2b; 1:1,000, MHC IIa; Alexa Fluor 488 IgG1; 1:1,000, and MHC IIb; Alexa Fluor 568 IgM; 1:1,000 for 1 hour at room temperature. Following incubation, slides were washed three times in PBS for 5-minutes per wash and mounted with ProLong antifade reagent (Life Technologies, Thermo Fisher Scientific).

Images were acquired after 24 hours through the AOMF facility at the University of Toronto. A total of 30–40 muscle fibers per fiber type were selected randomly throughout the cross section and traced with ImageJ software to assess cross-sectional area (CSA) and minimal-feret diameter (MFD). Muscle fibers exhibiting a black appearance were classified as MHC IIx.

### Statistics

Results are expressed as mean ± SD. The level of significance was established at *p* <0.05 for all statistics. Before statistical analyses, outliers were omitted in accordance with ROUT testing (Q = 0.5%), then tested for normality using D’Agostino–Pearson omnibus normality test. One-way ANOVAs were performed on normally distributed data, while Kruskal-Wallis test was performed on non-normally distributed data. Data with two independent variables that did not pass normality were first log transformed then analyzed using a standard two-way ANOVA. Post-hoc analyses were performed using the two-stage step-up method of Benjamini, Krieger, and Yekutieli to adjust for false discovery rate for data that demonstrated a significant interaction. While the results presented in **Fig.4** and **SFig.4** are non transformed data, the transformed data are shown in **SFig. 5** for reference. The objective of this study was to investigate the role of mitochondrial H_2_O_2_ in mediating apoptosis and necroptosis during cancer-induced atrophy. Thus, the following comparisons were not conducted: Early-EOC vs Late-EOC-SkQ1, and Early-EOC-SkQ1 and Late-EOC. The Mantel-Cox test was used to compute a curve comparison of survival data (p <0.05). All Statistical analyses were performed on Graphpad Prism 10 (La Jolla, CA, USA). Corresponding statistical tests are provided in figure legends.

## Results

### SkQ1 does not prevent ovarian cancer-induced body weight and fat loss, nor does it influence survivability

ID8 epithelial ovarian cancer (EOC) cells were injected directly beneath the ovarian bursa of syngeneic female mice and compared to PBS-injected sham mice (**Fig. 1A**). Half of the EOC-injected mice in both the early- and late-stage groups received the mitochondria targeting antioxidant SkQ1 in their drinking water. There were no significant differences in the survival rates between the Late-EOC-SkQ1 group and the normal drinking water group (**Fig. 1B**).

**Figure 1.**
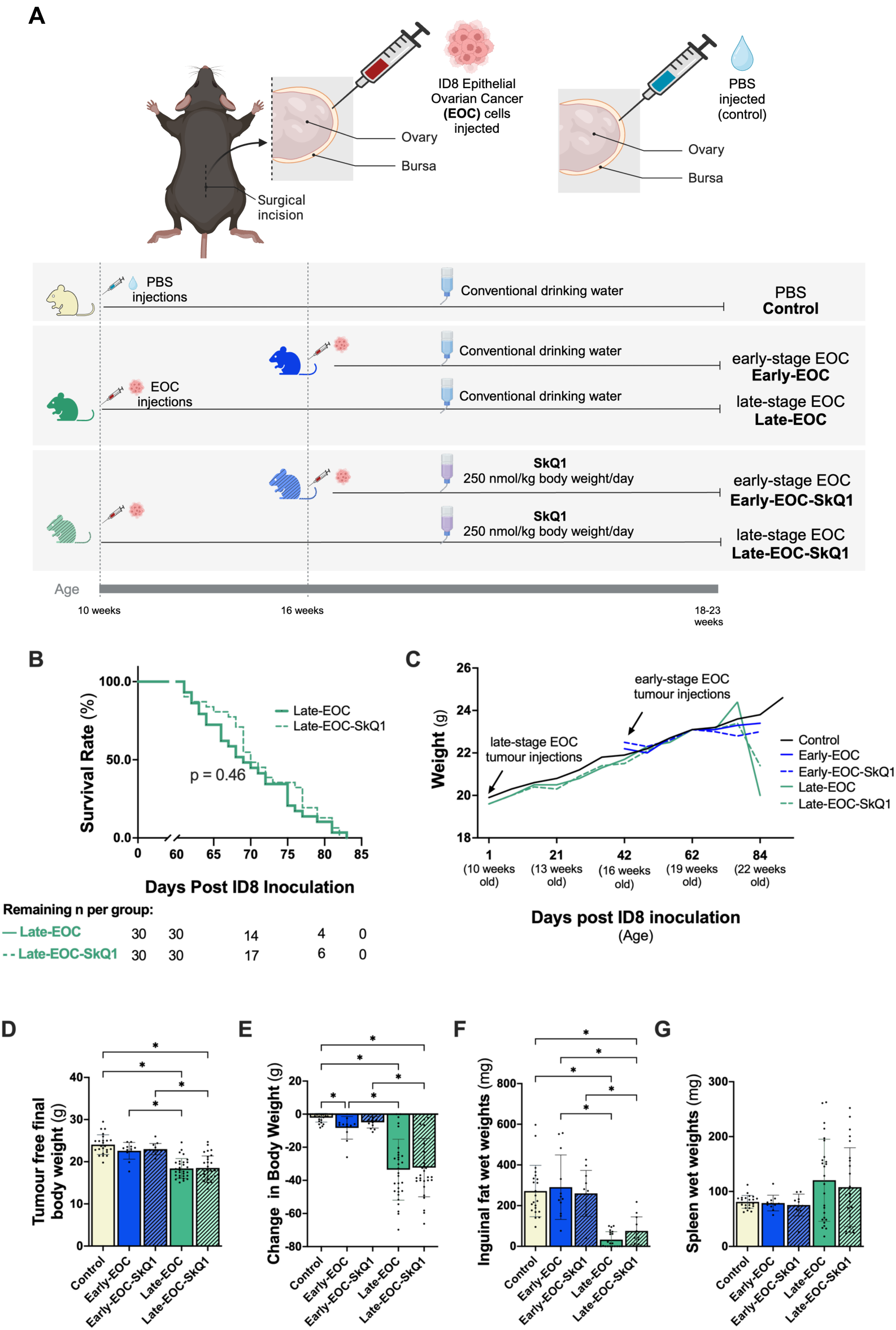
The effects of ovarian cancer progression and SkQ1 treatment on body weight, survival, and adipose tissue. **A)** Study design. Transformed murine ovarian surface (ID8) epithelial cells (EOC) from C57BL/6J mice were injected beneath the ovarian bursa of syngeneic adult female mice and compared to PBS-injected controls (PBS-control, n=24). Early–stage tumours progressed for 35-46 days (22-23 weeks of age) while Late-stage tumours progressed for 58-82 days (18-22 weeks of age). PBS Control mice were injected at 10 weeks of age and aged to 23 weeks (91-105 days post PBS injection). Following EOC injections, half the mice received the mitochondrial-targeting antioxidant, SkQ1 in drinking water (Early-EOC-SkQ1 n=12, Late-EOC-SkQ1 n=30), while the rest received standard water (Early-EOC n=12, Late-EOC n=30). **B)** Survival rate of Late-EOC and Late-EOC-SkQ1 groups. Late-stage EOC mice were aged to the latest timepoint possible prior to presentation of combinations of the following criteria: 10% body weight loss, > 20mL of ascitic volume collected during paracentesis, > 5 ascites paracentesis taps completed, and/or subjective changes in behavioural patterns consistent with removal criteria as per animal care guidelines (self-isolation, ruffled fur/poor self-grooming and irregular gait). During the ageing process, 6 Late-EOC mice and 4 Late-EOC-SkQ1 mice expired. **C)** Timeline of average bodyweight per group post tumour and PBS injections**. D)** Tumour free final body weight and **E)** difference between peak and final body weights per group (peak body weight – final body weight, n=12-30). **F)** Inguinal fat wet weights at endpoint (n=11-23). **G)** Spleen wet weights at endpoint (n=12-26). Survival curve analysis in **B** was performed using the Mantel-Cox survival test. Analyses for **D-G** were conducted using either a standard one-way ANOVA or Kruskal-Wallis test, depending on whether data was normally distributed. These were followed by a two-step step-up method of Benjamini, Krieger, and Yekutieli for post-hoc analysis. Results represent mean ± SD. **\*** = p < 0.05.

Late-EOC tumour-bearing mice exhibited reduced body weights and inguinal fat pads, with no significant changes due to SkQ1 treatment (**Fig. 1C-F**). Spleens were weighed at endpoint as an indicator of inflammation, but there were no differences between groups (**Fig. 1G**).

### SkQ1 does not affect tumour growth nor ascites fluid development

Primary ovarian tumours (at the original site of EOC injection) were excised and weighed at the endpoint. As expected, the primary tumours were significantly larger in both late-stage EOC groups. (**Fig. 2A**). Mice in both late-stage EOC groups developed ascites fluid, necessitating an average of 2.5 abdominal paracentesis procedures and approximately 10 mL of ascites fluid removed per group (**Fig. 2B-D**). SkQ1 did not induce any changes in tumour size, ascites fluid development or metastasis of EOC cells to the diaphragm (**SFig. 1**)

**Figure 2.**
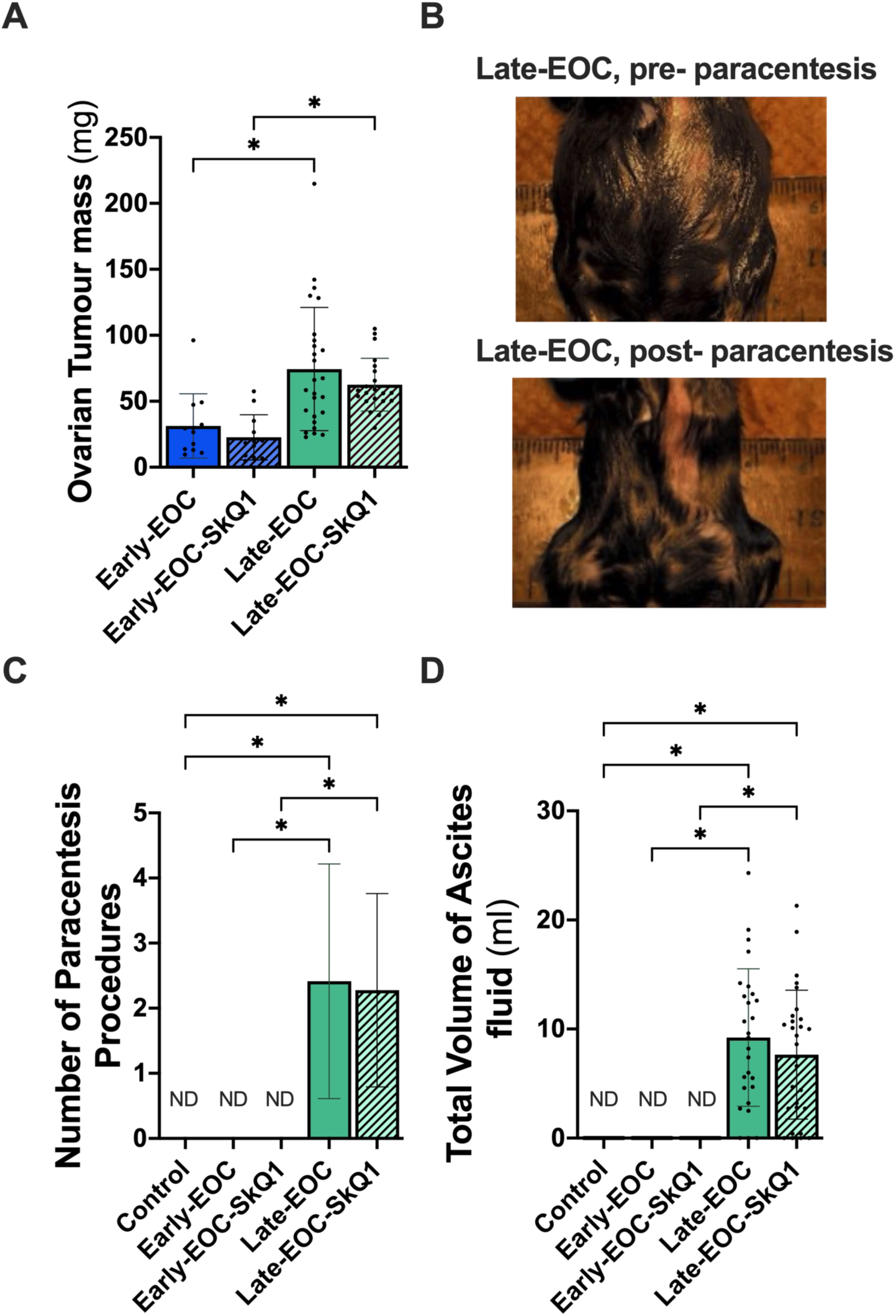
The effect of SkQ1 on primary tumour growth and ascites fluid development. **A)** Primary ovarian tumour mass at endpoint (n=12-26). **B)** Abdomen size prior to and post ascites fluid removal via paracentesis procedure. **C)** Average number of paracentesis procedures per mouse. **D)** Total volume of Ascites fluid removed per mouse (n= 12-26). **ND:** None detected, these groups did not develop ascites. Analyses for **A, C, and D** were conducted using either a standard one-way ANOVA or Kruskal-Wallis test, depending on whether data was normally distributed. These were followed by a two-step step-up method of Benjamini, Krieger, and Yekutieli for post-hoc analysis. Results represent mean ± SD. **\*** = p<0.05.

### SkQ1 does not prevent ovarian cancer-induced skeletal muscle atrophy

Hindlimb muscle wet weights were reduced in both late-stage untreated EOC group which was unaffected with treatment (**Fig. 3A**). The soleus muscle demonstrated a transient increase in wet weight in the untreated early-stage EOC group followed by a decrease in both late-stage EOC groups (**Fig. 3A**).

**Figure 3.**
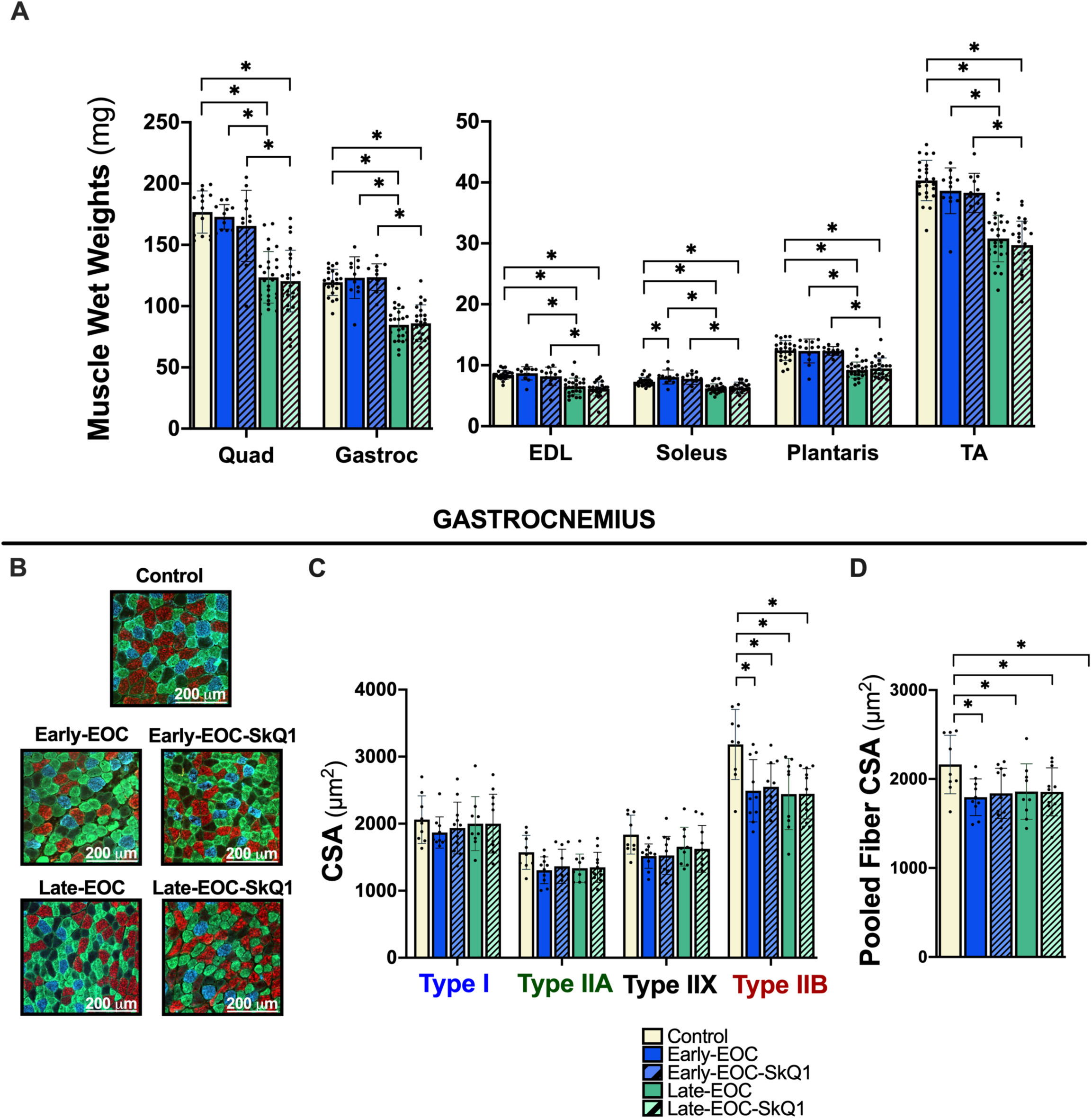
Evaluation of endpoint skeletal muscle wet weights and gastrocnemius fiber type atrophy with ovarian cancer progression. **A**) Hindlimb muscle wet weights at endpoint. **B)** Representative images of gastrocnemius cross-sections. n= 12 - 26. Blue corresponds to type I fibers, green: type IIA, black: type IIX, and red: type IIB fibers. **C)** Cross-sectional area (CSA) of gastrocnemius muscle fibers stained for myosin-heavy chain (MHC) isoforms, n=9 −12. **D)** Pooled gastrocnemius muscle fiber CSA, n=9 −12. **Quad** : quadriceps, **Gastroc** : Gastrocnemius, **EDL**: Extensor Digitorum Longus, **TA**: Tibialis Anterior. Analyses for **A, C and D** were conducted using either a standard one-way ANOVA or Kruskal-Wallis test, depending on whether data was normally distributed. These were followed by a two-step step-up method of Benjamini, Krieger, and Yekutieli for post-hoc analysis. Results represent mean ± SD. **\*** = p < 0.05.

The gastrocnemius muscle was stained for myosin heavy chain (MHC) isoforms to examine fiber-specific changes (**Fig. 3B**). Cross-sectional area (CSA) of type IIB fibers significantly decreased in all EOC groups, and this effect was maintained when all fiber types were pooled together, indicating that the average fiber size per group was lower in the EOC groups (**Fig. 3C and D**). Minimal Feret Diameter (MFD) was also measured as a second indicator of fiber-size changes. Similar to CSA, MFD of type IIB fibers decreased in all cancer groups (**SFig. 2A**). However, SkQ1 did not rescue gastrocnemius CSA or MFD in either the early- or late-stage groups. We also examined fibre-type specific CSA of the quadriceps muscles, which revealed lower type IIB CSA only in the late-stage group compared to control (**SFig. 3B-C**). However, MFD analysis did not reveal the same effect.

### SkQ1 ameliorated ovarian cancer-induced mitochondrial H_2_O_2_ emission in mixed gastrocnemius muscles of Late-EOC mice

Electron transport chain complex protein contents (specific subunits) were measured as an indirect indicator of mitochondrial quantity. Protein content of complexes I and II subunits decreased in the Late-EOC group compared to the Early-EOC group, and this decrease remained evident when protein contents of all complexes were pooled (**Fig. 4B**). However, these decreases in Complex I and II were not observed in the Late-EOC-SkQ1 group compared to the Early-EOC-SkQ1 group, suggesting that SkQ1 may have exerted a preservation effect on Complex I and II quantity during cancer progression.

**Figure 4.**
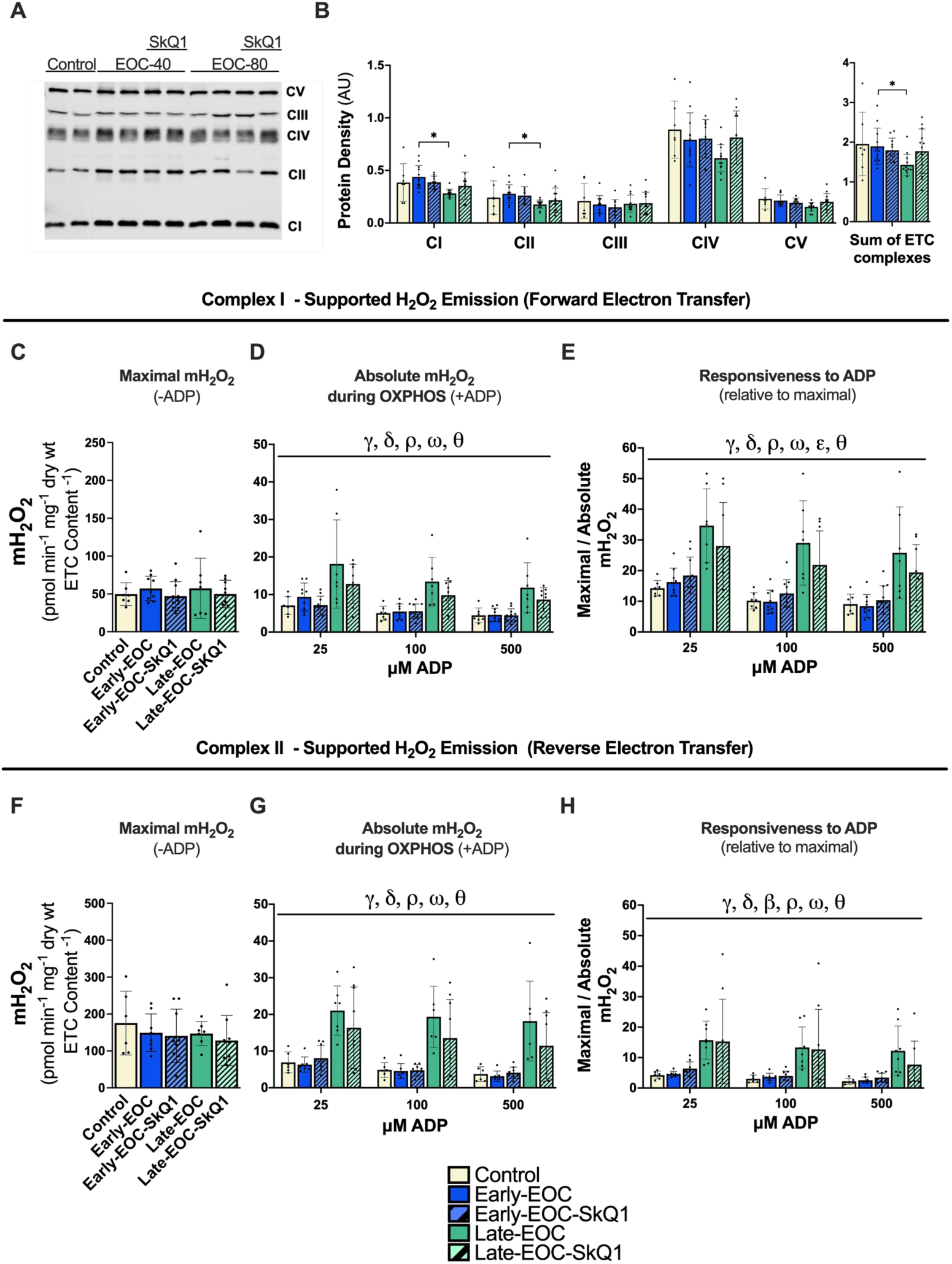
Changes in electron transport chain subunit content and Complex I- and II-supported H_2_O_2_ emission in white gastrocnemius of EOC-bearing mice subjected to either SkQ1 or normal drinking water. **A)** Representative western blot of electron transport chain (ETC) complex subunits in white gastrocnemius. **B)** Protein density of ETC subunits (see methods) in the white gastrocnemius, n=7-13. **C)** Complex I-supported (NADH supplied via pyruvate and malate) maximal mH_2_O_2_ (in the absence of ADP; - ADP) normalized to ETC content (n= 7-11). **D)** Absolute rates of Complex I -supported mH_2_O_2_ in the presence of ADP (+ADP) to model kinetics during oxidative phosphorylation normalized to ETC contents. **E)** mH_2_O_2_ during oxidative phosphorylation (+ADP)expressed relative to maximal (-ADP) rates as an index of mitochondrial responsiveness to ADP’s attenuating effects. **F)** Complex II-supported (FADH_2_ supplied via succinate) maximal mH_2_O_2_ (-ADP) normalized to ETC content (n= 7-11). **G)** Absolute rates of Complex II-supported mH_2_O_2_ emission in the presence of ADP (+ADP) to model kinetics during oxidative phosphorylation normalized to ETC content. **H)** mH_2_O_2_ during oxidative phosphorylation (+ADP) expressed relative to maximal (-ADP) rates as an index of mitochondrial responsiveness to ADP’s attenuating effects. **CI**: Complex I, **CII**: Complex II, **CIII**: Complex III, **CIV**: Complex IV, **CV**: Complex V, **ETC**: Electron Transport Chain, **H_2_O_2_**: Hydrogen Peroxide. Analyses for **B, C, F** were conducted using either a standard one-way ANOVA or Kruskal-Wallis test, depending on whether data was normally distributed. Analyses for **D**, **E**, **G** and **H** were conducted using a standard two-way ANOVA. These were followed by a two-step step-up method of Benjamini, Krieger, and Yekutieli for post-hoc analysis. Results represent mean ± SD. **β** = p <0.05 Control vs Early-EOC-SkQ1, **γ** = p < 0.05 Control vs Late-EOC, **δ** = < 0.05 Control vs Late-EOC-SkQ1, **ε** = p < 0.05 Early-EOC vs Early-EOC-SkQ1, **θ** = p < 0.05 Late-EOC vs Late-EOC-SkQ1, **ρ** = p < 0.05 Early-EOC vs Late-EOC, **ω** = Early-EOC-SkQ1 vs Late-EOC-SkQ1.

We then assessed mH_2_O_2_ using various substrates and inhibitors to identify changes in mH_2_O_2_ at specific sites. Complex I-supported (NADH supplied via pyruvate and malate) maximal mH_2_O_2_, in the absence of ADP-stimulation of oxidative phosphorylation, did not change with EOC or SkQ1 administration (**Fig. 4C, SFig. 4A**). ADP was then titrated to assess its ability to attenuate mH_2_O_2_ during oxidative phosphorylation (OXPHOS). Interestingly, mH_2_O_2_ was heightened in both late-stage EOC groups compared to control and their early-stage EOC counterparts, suggesting ADP is less effective at suppressing mH_2_O_2_ at Late-EOC (**SFig. 4B**). Notably, normalization of mH_2_O_2_ data to ETC contents revealed a significant decrease in Late-EOC-SkQ1 compared to Late-EOC, indicating that SkQ1 attenuated mH_2_O_2_ in late-stage cancer (**Fig. 4D**).

To further decipher whether the changes in mH_2_O_2_ during oxidative phosphorylation were due to mitochondrial responsiveness to ADP versus differences in initial maximal mH_2_O_2_, absolute rates in the presence of ADP were normalized to maximal mH_2_O_2_ (-ADP) and expressed as a percentage. The same pattern held, with both late-stage EOC groups exhibiting heightened mH_2_O_2_, which appeared to be attenuated by SkQ1 (**Fig. 4E**). A further pattern emerged regarding responsiveness to ADP, with Early-EOC-SkQ1 producing more H_2_O_2_ compared to Early-EOC. Collectively, these results indicated that changes in mH_2_O_2_ during cancer and in response to SkQ1 are due to a specific mechanism involving altered mitochondrial responsiveness to ADP.

Complex II-supported (FADH_2_ supplied via succinate) maximal and absolute mH_2_O_2_ during oxidative phosphorylation (via reverse electron flow) yielded the same results (**SFig. 4C-D, Fig. 4F-H**). mH_2_O_2_ from the pyruvate dehydrogenase complex (in the presence of pyruvate and rotenone to block Complex I oxidation of NADH and induce back-pressure on PDC) did not change with ovarian cancer progression or SkQ1 (**SFig. 4E**). Similarly, Complex III-stimulated mH_2_O_2_ (Antimycin A) also did not change with ovarian cancer progression or SkQ1 (**SFig. 4F**).

### Ovarian cancer progression leads to greater susceptibility to calcium-stress induced mPT and increased caspase-3 and caspase-9 activity in white gastrocnemius fibers

Mitochondrial permeability transition (mPT) was induced via calcium titrations in white gastrocnemius PmFB to assess the ability of mitochondria to withstand calcium stress (calcium retention capacity) and further their susceptibility to mPT. There were no differences in calcium retention capacity among any of the groups (**Fig. 5B**). However, the susceptibility of fibers to undergo mPT increased with ovarian cancer progression, with a greater number of fibers apparently undergoing mPT in both late-stage EOC groups compared to control, indicating that ovarian cancer makes WG fibers more prone to mPT (**Fig. 5C**).

**Figure 5.**
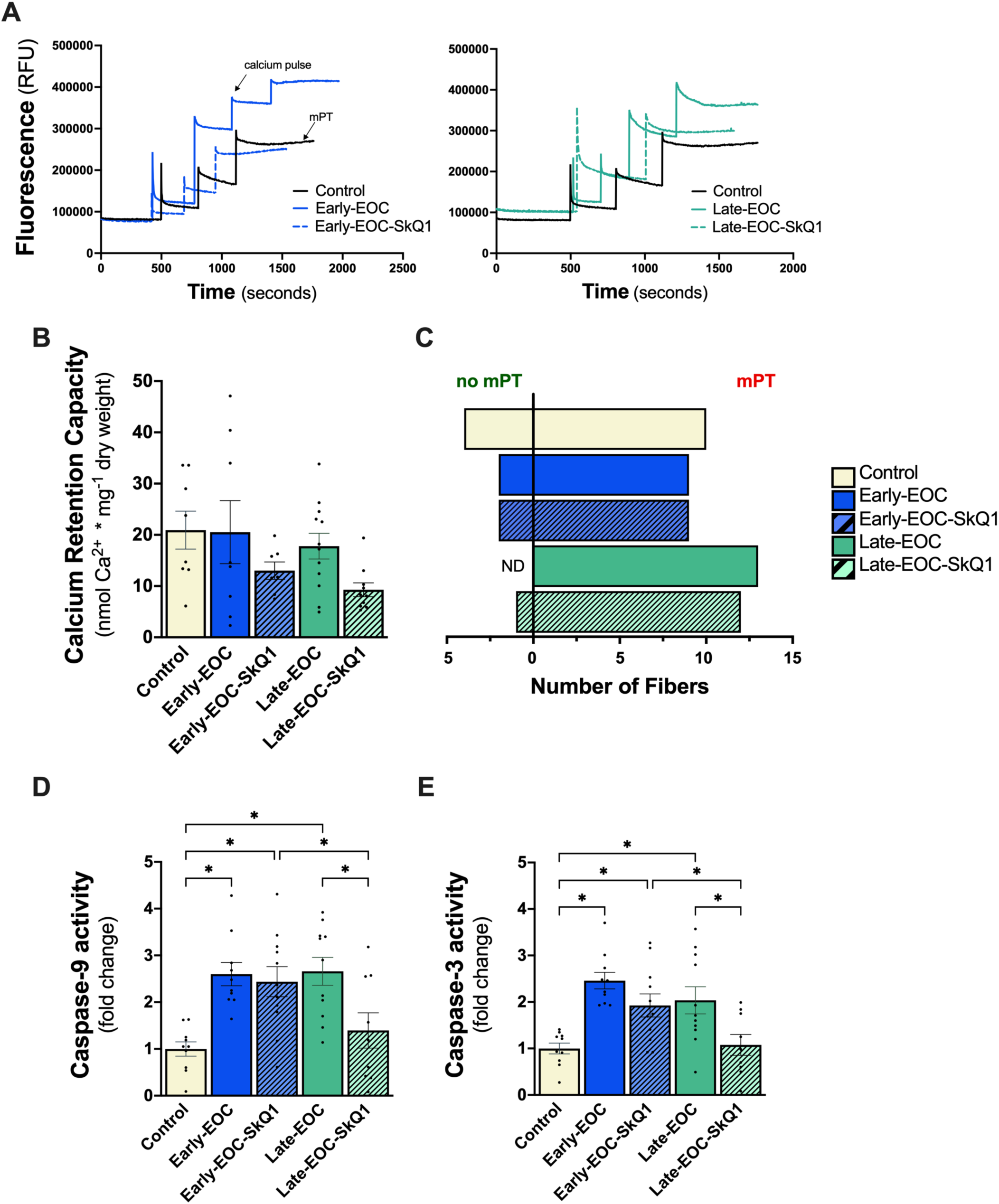
Calcium-stress induced mPT and caspase-9 and −3 activity in white gastrocnemius muscles of EOC-tumour bearing mice subjected to either SkQ1 or normal drinking water. **A)** Representative traces of calcium titrations and induction of mitochondrial permeability transition (mPT) in white gastrocnemius muscles. **B)** Calcium retention capacity required to initiate mPT in white gastrocnemius fibers, (n =9-13). **C)** Incidence of calcium-stress induced mPT in mixed gastrocnemius fibers (right side) vs fibers that did not exhibit mPT (left side), (n=9-13). **D)** Fold change in caspase-9- and **E)** caspase-3 activity, n=9-11. **ND**: none detected. Analyses for **B, D, and E** were conducted using either a standard one-way ANOVA or Kruskal-Wallis test, depending on whether data was normally distributed. These were followed by a two-step step-up method of Benjamini, Krieger, and Yekutieli for post-hoc analysis. Results represent mean ± SD. **\*** = p < 0.05.

Caspase-9 and −3 activity increased 2.5-fold in both early-stage EOC groups and remained elevated in the Late-EOC group (**Fig. 5D-E**). Notably, in the Late-EOC-SkQ1 group, caspase-9 and −3 activity decreased reaching levels comparable to Control caspase activity. This suggests that SkQ1 helps mitigate the increased caspase activity associated with EOC progression.

### The capacity for white gastrocnemius muscle to undergo necroptosis is heightened during early ovarian cancer progression

Protein content of total RIPK1, RIPK3, and MLKL, as well as their phosphorylated counterparts, were measured as markers of necroptosis. First, the accuracy of necroptosis antibodies was assessed using positive (C2C12 model of necroptosis, DMD tissue) and negative (RIPK3 KO tissue) controls (**SFig. 6-8**).

**Figure 6.**
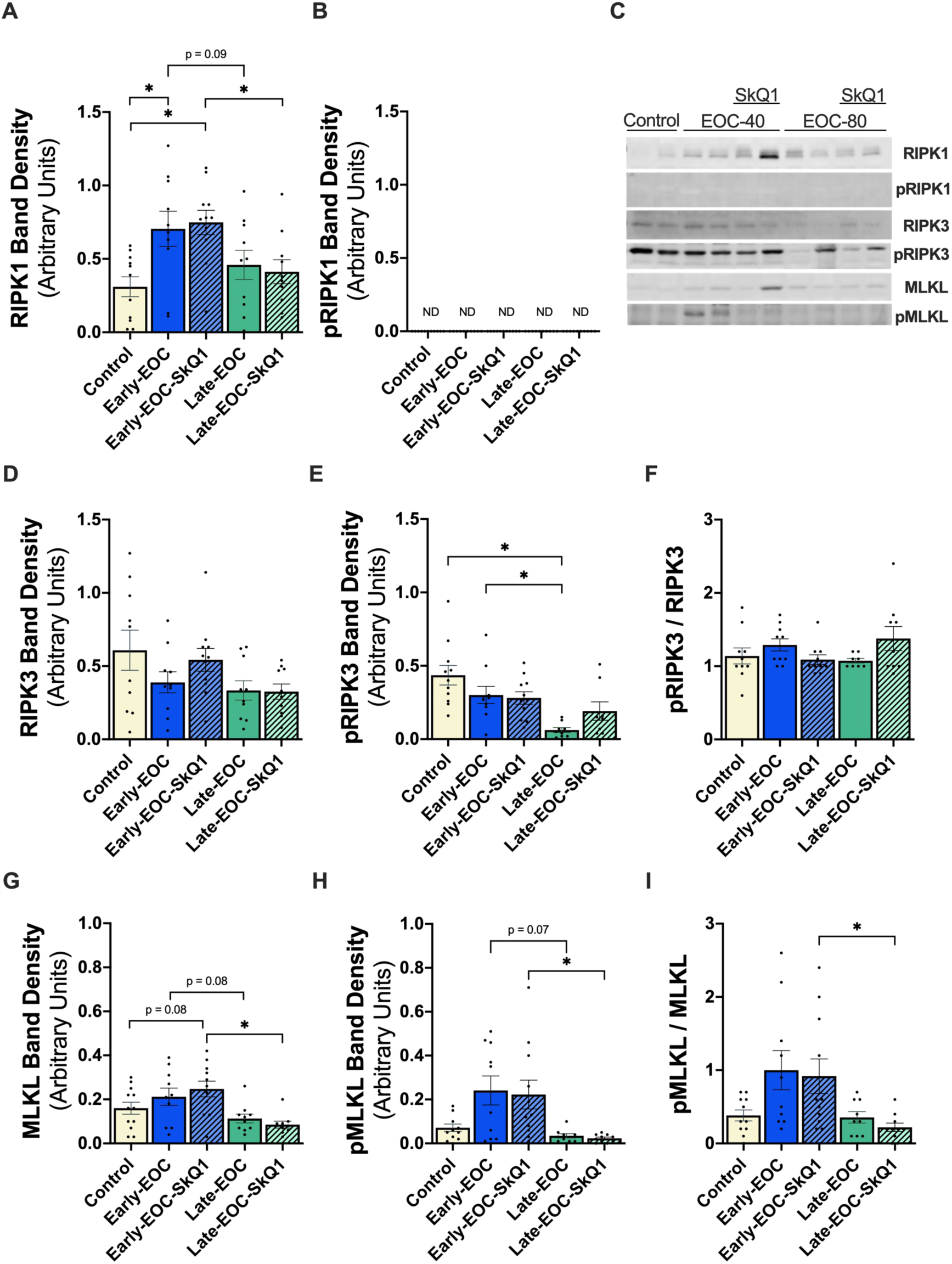
Assessment of necroptosis markers in white gastrocnemius muscles of EOC-tumour bearing mice subjected to either SkQ1 or normal drinking water. **A)** Total RIPK1 and **B)** phosphorylated-RIPK1 protein content in the white gastrocnemius muscle (n=10-12). **C)** Representative western blot of RIPK1, RIPK3, MLKL and their phosphorylated counterparts in white gastrocnemius muscle. **D)** Total RIPK3 and **E)** phosphorylated RIPK3 protein content in white gastrocnemius**. F)** phosphorylated RIPK3 normalized to total RIPK3 protein content**. G)** Total MLKL and **H)** phosphorylated protein content in white gastrocnemius muscle. **I)** phosphorylated MLKL normalized to total MLKL protein content. **RIPK1**: Receptor Interacting Serine/Threonine Protein Kinase 1, **RIPK3**: Receptor Interacting Serine/Threonine Protein Kinase 3, **MLKL**: Mixed-Lineage Kinase Domain-Like Protein, **pRIPK1**: phosphorylated RIPK1, **pRIPK3**: phosphorylated RIPK3, **pMLKL**: phosphorylated MLKL. **ND**: none detected. Analyses for **A**, **B**, **D**, **E**, **F**, **G** and **H** were conducted using either a standard one-way ANOVA or Kruskal-Wallis test, depending on whether data was normally distributed. These were followed by a two-step step-up method of Benjamini, Krieger, and Yekutieli for post-hoc analysis. Results represent mean ± SD. **\*** = p < 0.05.

Total RIPK1 protein content increased in both early-stage EOC groups compared to control, that was not maintained in both late-stage EOC groups (**Fig. 6A**). Assessment of phosphorylated RIPK1 did not yield detectable bands in any group, leaving uncertainty about whether the increased RIPK1 total protein at Early-EOC corresponds to heightened activation (via phosphorylation) (**Fig. 6B**).

Divergent results were observed for RIPK3 and MLKL compared to RIPK1. Total RIPK3 showed no changes among any of the groups (**Fig. 6D**). Phosphorylated RIPK3 decreased in the Late-EOC group compared to control and Early-EOC, but this effect was lost when normalized to total RIPK3 content, indicating no significant differences among the groups (**Fig. 6E-F**).

For MLKL, both total and phosphorylated protein content decreased in the Late-EOC-SkQ1 group compared to the Early-EOC-SkQ1 group, though no differences were observed when compared to the control group (**Fig. 6G-H**). This decrease persisted when phosphorylated MLKL was normalized to total MLKL content, suggesting a consistent reduction in MLKL activation in the Late-EOC-SkQ1 group (**Fig. 6I**).

## Discussion

The degree to which mitochondrial-linked cell death pathways contribute to muscle atrophy in cancer remains unclear. Recent discoveries that increased mitochondrial reactive oxygen species manifests early during cancer cachexia suggests mitochondrial-linked cell death pathways like apoptosis and necroptosis could contribute to atrophy during cancer. In this study, we investigated whether mitochondrial H_2_O_2_-mediated cell death contributes to muscle atrophy during cancer progression and whether reducing mH_2_O_2_ with the well-established mitochondrial-targeted antioxidant SkQ1 could prevent cell death and preserve muscle mass. Using a robust orthotopic model of metastatic ovarian cancer in adult immunocompetent mice, we show for the first time that mitochondrial H_2_O_2_ increases at late-stage ovarian cancer (stage III) in type IIb rich gastrocnemius muscle fibres, which does not directly drive muscle atrophy through mitochondrial-linked apoptosis or necroptosis. Importantly, our study provides novel evidence that mitochondrial H_2_O_2_, apoptosis, and necroptosis are not major contributors to muscle atrophy in the type II B-rich white gastrocnemius during ovarian cancer progression (**Fig. 7**). These discoveries contrast previous reports of mitochondrial-targeted antioxidants preventing atrophy during cancer. Collectively, these results underscore the need for further research to determine whether mitochondrial contributions to atrophy differ across muscle groups or type of cancer.

**Figure 7.**
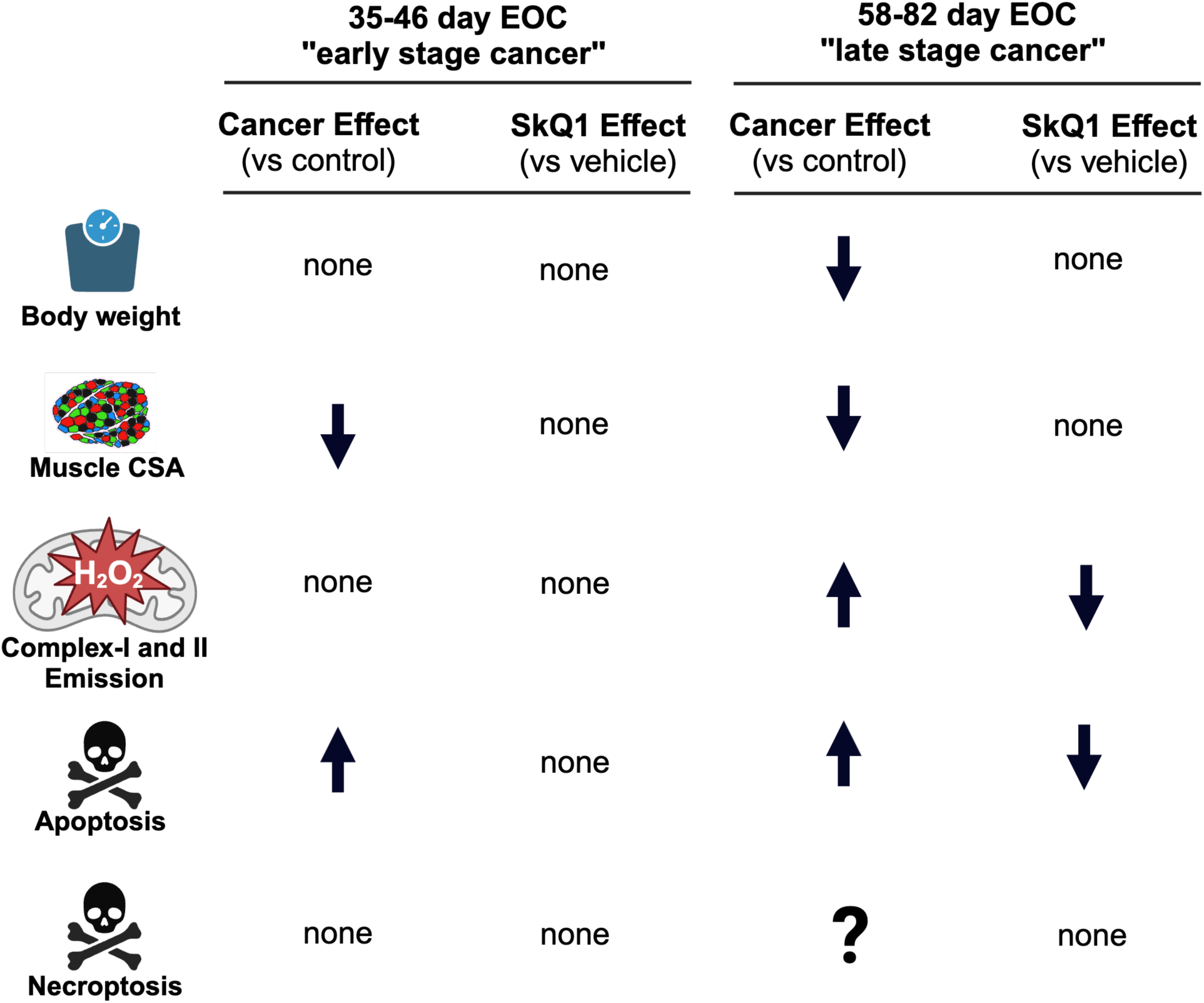
Summary of findings in the gastrocnemius muscles. “↑” represents increase, “↓” represents decrease, “?” represents inconclusive findings, none represents no statistically relevant changes in either EOC groups compared to control or SkQ1 groups compared to EOC groups at the given stage. Muscle cross-sectional area (CSA) data obtained from the mixed gastrocnemius while complex I and II mH_2_O_2_, apoptosis and necroptosis markers data were obtained using white gastrocnemius muscles.

### SkQ1 does not rescue EOC-induced skeletal muscle atrophy

Several mouse models of cancer cachexia exhibit elevated mitochondrial ROS prior to atrophy (Brown *et al*., 2017; Delfinis *et al*., 2022) which corresponds to a pre-metastatic early stage of cancer. However, the degree to which mitochondrial ROS contribute to skeletal muscle atrophy has been unclear due to discrepant findings in the literature (Brown *et al*., 2017; Halle *et al*., 2019; Delfinis *et al*., 2022). Our findings of no relationship between mitochondrial H_2_O_2_ and atrophy are consistent with previous reports that mice over-expressing the H_2_O_2_-scavenging antioxidant catalase in the mitochondria are not protected from breast cancer-induced muscle atrophy in the soleus (Gilliam *et al*., 2016). However, the cardiolipin-targeting peptide SS-31 prevented increases in mitochondrial ROS and loss of diaphragm and EDL fibre cross sectional area in C26 cancer mice (Smuder *et al*., 2020). Additionally, the mitochondrial ubiquinone-based antioxidant MitoQ, which is thought to be similar in mechanism to the plastoquinone component of SkQ1, prevented atrophy in muscle cell culture and partially prevented a loss of mass in TA and gastrocnemius of C26 mice but did not prevent a loss of fibre cross sectional area in the TA (Pin *et al*., 2022). Our group recently reported that SkQ1 prevented atrophy in male gastrocnemius but seemed to induce atrophy in female plantaris during colorectal (C26) cancer (Tsitkanou *et al*., 2024). Therefore, our findings align with an emerging pattern in the literature that points to a complex relationship between mitochondrial ROS, mitochondrial therapeutics, and muscle atrophy during cancer. This body of evidence underscores the need for careful in-depth consideration of the mechanisms of action of mitochondrial-therapeutics (plastoquinone/quinone vs cardiolipin-targeting small peptide therapeutics) and whether the role of mitochondrial bioenergetics in mediating atrophy differs across types or duration of cancer, muscle groups, or biological sex, and whether methods of assessing atrophy (weights vs cross-sectional area) influence interpretations. This pattern cannot be explained by existing understanding of the effects of mitochondrial ROS on atrophy which highlights the opportunity for asking new questions regarding the precise role of mitochondria in remodeling muscle during cancer.

### Mitochondrial-induced apoptosis is elevated in skeletal muscle during ovarian cancer and is prevented by SkQ1

Although multiple pathways can initiate apoptosis, this study focused specifically on mitochondrial-induced apoptosis by exploring the relationship between mPT and caspase-9 and caspase-3 activity. mPT is known to trigger the formation of the ‘apoptosome’, comprising APAF-1, cytochrome *c* and procaspase-9 (Bock & Tait, 2019; Sever *et al*., 2023). Once established, the apoptosome cleaves initiator procaspase-9, into its active form, caspase-9, which then cleaves and activates executioner caspase-3 (Bhola *et al*., 2009; Bock & Tait, 2019; Skinner *et al*., 2021; Sever *et al*., 2023). Classically, executioner caspases like caspase-3 then cleave key structural and functional proteins (Du *et al*., 2004; Bock & Tait, 2019; Skinner *et al*., 2021). Notably, caspase-3 has been demonstrated to cleave myofibrillar proteins in rat psoas muscle lysates *in vitro* (Du *et al*., 2004). As such, measures of caspase-9 and −3 activity are useful for interpreting mitochondrial-induced apoptosis.

Our discovery that ovarian cancer increases caspase-9 and −3 activity in white gastrocnemius during early- and late-stage ovarian cancer suggests an early role for mitochondrial-induced apoptosis. These results partially align with our mPT findings which showed a greater number of samples demonstrating calcium-induced mPT in late-stage but not early-stage. The reason for a lack of relationships between early stage mPT and caspase-9 and −3 activities is not clear but may be due to methodological considerations. While we used an established assay for assessing mPT, the findings can only be interpreted under conditions of calcium stress and as a ‘capacity’ measure (maximal calcium required to trigger mPT). It is possible that mPT occurred in vivo without changes in capacity for calcium-triggered mPT, but measuring such possibilities is not technically feasible with small muscle samples. Also, other triggers of mPT may not be captured in this assay. For example, increases in ROS or extreme reductions in membrane potential are also known to initiate mPT. These stimuli were not intentionally controlled during our assay, which means that future studies could employ different approaches to assess mPT in muscle during cancer. Likewise, mPT can also be assessed with time-to-opening in response to calcium challenges if a single large bolus is used rather than a titration of calcium as used in the present study which may reveal different insight into mPT dynamics. Nonetheless, our results support a role of calcium stress-induced mPT and apoptotic caspase-9 and −3 activity in late-stage ovarian cancer in atrophied white gastrocnemius muscle.

Remarkably, both atrophy and the elevated caspase-9 and −3 activity during early-stage cancer were detected before any observable changes in mitochondrial H_2_O_2_ emission or the likelihood of mPT. This suggests that caspase-9 and −3 may contribute to ovarian cancer-induced atrophy independently of mH_2_O_2_ emission or mPT, at least during early-stage cancer. In contrast, late-stage cancer demonstrated both increased mH_2_O_2_ and caspase-9 and −3 activity.

Notably, SkQ1 completely prevented caspase-9 and −3 activity at late-stage ovarian cancer but there was no apparent reduction in mPT in terms of the number of samples that demonstrated an opening. Nonetheless, the findings demonstrate that SkQ1 is capable of completely preventing mitochondrial-induced apoptosis during ovarian cancer concurrent with lower mH_2_O_2_. This is the first demonstration of a direct relationship between mH_2_O_2_ and apoptosis during cancer-induced atrophy. However, as discussed above, these relationships had no bearing on atrophy itself given SkQ1 did not prevent reductions in fibre cross sectional area or mass in the gastrocnemius as noted above. Therefore, these discoveries suggest that mitochondrial-induced apoptotic signaling was not a contributor to atrophy during ovarian cancer. The finding raises new questions regarding the precise role of caspase-9 and −3 during cancer and may lead to new avenues of research exploring how protein degradation may influence other properties of muscle fibres beyond their size.

Likewise, the degree to which catabolic processes are countered by anabolic responses to maintain fibre size is not clear.

### Inconsistent markers of necroptosis during ovarian cancer

In many cell types, classical necroptosis is dependent on the formation of the ‘necrosome’, a complex composed of RIPK1 and RIPK3, which undergo phosphorylation to drive the subsequent phosphorylation of MLKL monomers (Huang *et al*., 2013; Gan *et al*., 2019; Khoury *et al*., 2020; Chu *et al*., 2021; Oikonomou *et al*., 2021). MLKL monomers then oligomerize to form pores in the plasma membrane, leading to cellular leakage and death (Rodriguez *et al*., 2016; Samson *et al*., 2020). Mitochondria also play a pivotal role in this pathway, with mH_2_O_2_ oxidizing cysteine residues on RIPK1, thereby inducing its autophosphorylation and promoting further necrosome formation(Zhang *et al*., 2017).

In early-stage ovarian cancer, there is a notable increase in RIPK1 content in white gastrocnemius which coincides with a reduction in the cross-sectional area of these muscle fibers, suggesting a potential link between elevated RIPK1 levels and early muscle atrophy. However, phosphorylation of RIPK1 was not detected in any group despite preliminary validations of the antibody used within this study as shown in Supplemental data (**SFig. 6-8**). It is unclear whether RIPK1 phosphorylation is a highly transient event that may occur at other timepoints during cancer. Nonetheless, the increase in total RIPK1 content suggests that the capacity for necroptosis is increased in muscle during early-stage ovarian cancer. Thereby, further research into the potential influence of necroptosis on muscle atrophy during ovarian cancer is warranted which may involve inducing cancer in RIPK1 kinase-inactive models (Polykratis *et al*., 2014). Notably, RIPK1 is also a critical regulator upstream of various regulated cell death pathways, including extrinsic apoptosis and pyroptosis which were not assessed in this study (Galluzzi *et al*., 2018; Zhang *et al*., 2019; Yow *et al*., 2024).

RIPK3 and MLKL are downstream of RIPK1 and specific to necroptosis. However, despite the early increase in the potential for RIPK1-dependent necroptosis, phosphorylation of MLKL at late-stage cancer was comparable to controls, while phosphorylation of RIPK3 decreased below control levels. Further, this decline in RIPK3 phosphorylation was lost when normalized to total RIPK3 content. In this regard, the finding that phosphorylated RIPK3 decreased in late-stage cancer below control levels is difficult to interpret but may lead to new questions regarding potential compensatory attempts of muscle fibres to downregulate necroptosis even further below basal levels in late-stage cancer.

Phosphorylated MLKL was variable throughout cancer. While there was no difference between control and early-stage cancer, there was a significant decrease in late-stage vs early-stage groups. These findings might suggest a high degree of variability in this marker during ovarian cancer. Despite the elevated mH_2_O_2_ observed in late-stage ovarian cancer, there was no corresponding increase in RIPK3 or MLKL phosphorylation. This finding is somewhat unexpected given mitochondrial ROS has been shown to induce necroptosis (Zhang et al., 2017; Yang et al., 2018).

### Mitochondrial adaptation to SKQ1 administration with EOC development

While SkQ1 did not prevent atrophy in the white gastrocnemius, there is still opportunity to determine whether it can prevent atrophy in other muscle types or prevent loss of muscle strength which was not assessed in this study given the objective focused atrophy as a primary outcome. In this way, understanding the mechanism by which SkQ1 lowered mH_2_O_2_ could be considered when designing such investigations. SkQ1 is comprised of plastoquinone tethered to a lipophilic cationic chaperone (triphenylphosphonium, or TPP+) (Skulachev et al., 2023). The latter properties increase the potential for tissue penetration (lipophilic) and attraction to the relatively negatively charged mitochondrial matrix. At certain dosages, the plastoquinone component is believed to accept electrons from Complex I and III, particularly when electrons may be prone to transfer onto oxygen prematurely to generate superoxide prior to it’s dismutation to H_2_O_2_ (Ježek et al., 2017). While we did not see changes in Complex III-stimulated mH_2_O_2_ using Antimycin A, we found mH_2_O_2_ stimulated by reverse electron transfer from Complex II to I (succinate supported) was attenuated by SkQ1 in the late-stage EOC group which could be consistent with it’s known actions on Complex I. However, there was no effect of SkQ1 in maximal mH_2_O_2_ with either forward or reverse electron transfer to Complex I. Rather, the attenuating effect of SkQ1 was seen only in the presence of ADP. It is therefore possible that SkQ1 improved mitochondrial responsiveness to ADP attenuation of ROS production separate from a potential action on Complex I itself. It may be possible that chronic administration of SkQ1 could prevent ROS-inhibition of other components of the phosphorylation system regulating adenylate cycling, but this would require targeted assessments of proteins such as ANT and VDAC to be certain. Such potential indirect effects of the drug over chronic treatment were considered for other ROS-producing pathways including pyruvate dehydrogenase complex, but no effects were observed.

### Perspectives and Conclusion

SkQ1 treatment did not affect primary tumour size, ascites development, metastasis, or overall survival, increasing our confidence that SkQ1 does not promote cancer growth or exacerbate its effects in this model. The findings from this study contrast recent findings by our group that SkQ1 prevented atrophy in gastrocnemius of male mice during C26 colorectal cancer but promoted atrophy in plantaris of female mice. These findings underscore the potential to explore SkQ1 and other mitochondrial therapeutics in specific biological contexts of cancer cachexia (time, biological sex, muscle type, cancer type, etc).

In conclusion, SkQ1 prevented mitochondrial-induced apoptosis but not atrophy in gastrocnemius during ovarian cancer. This evidence supports a mechanistic role of mitochondrial ROS, likely in the form of H_2_O_2_, and mPT in triggering apoptotic caspase-9 and −3 activity in white gastrocnemius during cancer but does not support a role of this form of apoptosis in mediating atrophy itself. Limited changes in markers of necroptosis suggest a limited role of this pathway in mediating atrophy, although future studies could examine their potential role with models that up- or down-regulate key regulatory necroptotic proteins. These findings add to the growing body of literature of complex relationships between mitochondrial targeted therapeutics and muscle outcomes during ovarian cancer and highlight the need for carefully considering numerous biological and methodological factors in experimental design.

## Data availability

data are available upon reasonable request submitted to the corresponding author.

## References

Adhihetty PJ, O’Leary MFN, Chabi B, Wicks KL & Hood DA (2007). Effect of denervation on mitochondrially mediated apoptosis in skeletal muscle. J Appl Physiol 102, 1143–1151.

Anisimov VN, Egorov M V., Krasilshchikova MS, Lyamzaev KG, Manskikh VN, Moshkin MP, Novikov EA, Popovich IG, Rogovin KA, Shabalina IG, Shekarova ON, Skulachev M V., Titova T V., Vygodin VA, Vyssokikh MY, Yurova MN, Zabezhinsky MA & Skulachev VP (2011). Effects of the mitochondria-targeted antioxidant SkQ1 on lifespan of rodents. Aging (Albany NY) 3, 1110.

Ballarò R, Lopalco P, Audrito V, Beltrà M, Pin F, Angelini R, Costelli P, Corcelli A, Bonetto A, Szeto HH, O’connell TM & Penna F (2021). Targeting Mitochondria by SS-31 Ameliorates the Whole Body Energy Status in Cancer- and Chemotherapy-Induced Cachexia. Cancers 2021, Vol 13, Page 850 13, 850.

Baltgalvis KA, Berger FG, Peña MMO, Davis JM, White JP & Carson JA (2010). Activity level, apoptosis, and development of cachexia in Apc Min/+ mice. J Appl Physiol 109, 1155– 1161.

Belizário JE, Lorite MJ & Tisdale MJ (2001). Cleavage of caspases-1, −3, −6, −8 and −9 substrates by proteases in skeletal muscles from mice undergoing cancer cachexia. British Journal of Cancer 2001 84:8 84, 1135–1140.

Bellissimo CA, Delfinis LJ, Hughes MC, Turnbull PC, Gandhi S, DiBenedetto SN, Rahman FA, Tadi P, Amaral CA, Dehghani A, Cobley JN, Quadrilatero J, Schlattner U & Perry CGR (2023). Mitochondrial creatine sensitivity is lost in the D2.mdx model of Duchenne muscular dystrophy and rescued by the mitochondrial-enhancing compound Olesoxime. Am J Physiol Cell Physiol 324, C1141–C1157.

Bernardi P, Gerle C, Halestrap AP, Jonas EA, Karch J, Mnatsakanyan N, Pavlov E, Sheu SS & Soukas AA (2023). Identity, structure, and function of the mitochondrial permeability transition pore: controversies, consensus, recent advances, and future directions. Cell Death & Differentiation 2023 30:8 30, 1869–1885.

Bhola PD, Mattheyses AL & Simon SM (2009). Spatial and temporal dynamics of mitochondrial membrane permeability waves during apoptosis. Biophys J 97, 2222–2231.

Bock FJ & Tait SWG (2019). Mitochondria as multifaceted regulators of cell death. Nature Reviews Molecular Cell Biology 2019 21:2 21, 85–100.

Bonora M, Giorgi C & Pinton P (2021). Molecular mechanisms and consequences of mitochondrial permeability transition. Nature Reviews Molecular Cell Biology 2021 23:4 23, 266–285.

Brown JL, Lawrence MM, Ahn B, Kneis P, Piekarz KM, Qaisar R, Ranjit R, Bian J, Pharaoh G, Brown C, Peelor FF, Kinter MT, Miller BF, Richardson A & Van Remmen H (2020). Cancer cachexia in a mouse model of oxidative stress. J Cachexia Sarcopenia Muscle 11, 1688.

Brown JL, Rosa-Caldwell ME, Lee DE, Blackwell TA, Brown LA, Perry RA, Haynie WS, Hardee JP, Carson JA, Wiggs MP, Washington TA & Greene NP (2017). Mitochondrial degeneration precedes the development of muscle atrophy in progression of cancer cachexia in tumour-bearing mice. J Cachexia Sarcopenia Muscle 8, 926.

Busquets S, Deans C, Figueras M, Moore-Carrasco R, López-Soriano FJ, Fearon KCH & Argilés JM (2007). Apoptosis is present in skeletal muscle of cachectic gastro-intestinal cancer patients. Clinical Nutrition 26, 614–618.

de Castro GS, Simoes E, Lima JDCC, Ortiz-Silva M, Festuccia WT, Tokeshi F, Alcântara PS, Otoch JP, Coletti D & Seelaender M (2019). Human Cachexia Induces Changes in Mitochondria, Autophagy and Apoptosis in the Skeletal Muscle. Cancers (Basel); DOI: 10.3390/CANCERS11091264.

Chu Q, Gu X, Zheng Q, Wang J & Zhu H (2021). Mitochondrial Mechanisms of Apoptosis and Necroptosis in Liver Diseases. Analytical Cellular Pathology; DOI: 10.1155/2021/8900122.

Delfinis LJ, Bellissimo CA, Gandhi S, DiBenedetto SN, Garibotti MC, Thuhan AK, Tsitkanou S, Rosa-Caldwell ME, Rahman FA, Cheng AJ, Wiggs MP, Schlattner U, Quadrilatero J, Greene NP & Perry CGR (2022). Muscle weakness precedes atrophy during cancer cachexia and is linked to muscle-specific mitochondrial stress. JCI Insight; DOI: 10.1172/JCI.INSIGHT.155147.

Delfinis LJ, Ogilvie LM, Khajehzadehshoushtar S, Gandhi S, Garibotti MC, Thuhan AK, Matuszewska K, Pereira M, Jones RG, Cheng AJ, Hawke TJ, Greene NP, Murach KA, Simpson JA, Petrik J & Perry CGR (2024). Muscle weakness and mitochondrial stress occur before severe metastasis in a novel mouse model of ovarian cancer cachexia. Mol Metab 86, 101976.

Du J, Wang X, Miereles C, Bailey JL, Debigare R, Zheng B, Price SR & Mitch WE (2004). Activation of caspase-3 is an initial step triggering accelerated muscle proteolysis in catabolic conditions. J Clin Invest 113, 115.

Fearon K, Strasser F, Anker SD, Bosaeus I, Bruera E, Fainsinger RL, Jatoi A, Loprinzi C, MacDonald N, Mantovani G, Davis M, Muscaritoli M, Ottery F, Radbruch L, Ravasco P, Walsh D, Wilcock A, Kaasa S & Baracos VE (2011). Definition and classification of cancer cachexia: an international consensus. Lancet Oncol 12, 489–495.

Fedorov A V., Chelombitko MA, Chernyavskij DA, Galkin II, Pletjushkina OY, Vasilieva T V., Zinovkin RA & Chernyak B V. (2022). Mitochondria-Targeted Antioxidant SkQ1 Prevents the Development of Experimental Colitis in Mice and Impairment of the Barrier Function of the Intestinal Epithelium. Cells; DOI: 10.3390/CELLS11213441.

Galluzzi L et al. (2018). Molecular mechanisms of cell death: recommendations of the Nomenclature Committee on Cell Death 2018. Cell Death & Differentiation 2018 25:3 25, 486–541.

Gan I, Jiang J, Lian D, Huang X, Fuhrmann B, Liu W, Haig A, Jevnikar AM & Zhang ZX (2019). Mitochondrial permeability regulates cardiac endothelial cell necroptosis and cardiac allograft rejection. American Journal of Transplantation 19, 686–698.

Gilliam LAA, Lark DS, Reese LR, Torres MJ, Ryan TE, Lin C Te, Cathey BL & Darrell Neufer P (2016). Targeted overexpression of mitochondrial catalase protects against cancer chemotherapy-induced skeletal muscle dysfunction. Am J Physiol Endocrinol Metab 311, E293–E301.

Glass DJ (2010). A Critique of the Hypothesis, and a Defense of the Question, as a Framework for Experimentation. Clin Chem 56, 1080–1085.

Greenaway J, Henkin J, Lawler J, Moorehead R & Petrik J (2009). ABT-510 induces tumor cell apoptosis and inhibits ovarian tumor growth in an orthotopic, syngeneic model of epithelial ovarian cancer. Mol Cancer Ther 8, 64.

Greenaway J, Moorehead R, Shaw P & Petrik J (2008). Epithelial-stromal interaction increases cell proliferation, survival and tumorigenicity in a mouse model of human epithelial ovarian cancer. Gynecol Oncol 108, 385–394.

Halle JL, Pena GS, Paez HG, Castro AJ, Rossiter HB, Visavadiya NP, Whitehurst MA & Khamoui A V. (2019). Tissue-specific dysregulation of mitochondrial respiratory capacity and coupling control in colon-26 tumor-induced cachexia. Am J Physiol Regul Integr Comp Physiol 317, R68–R82.

Huang CY, Kuo WT, Huang YC, Lee TC & Yu LCH (2013). Resistance to hypoxia-induced necroptosis is conferred by glycolytic pyruvate scavenging of mitochondrial superoxide in colorectal cancer cells. Cell Death & Disease 2013 4:5 4, e622–e622.

Hughes MC, Ramos S V., Turnbull PC, Rebalka IA, Cao A, Monaco CMF, Varah NE, Edgett BA, Huber JS, Tadi P, Delfinis LJ, Schlattner U, Simpson JA, Hawke TJ & Perry CGR (2019). Early myopathy in Duchenne muscular dystrophy is associated with elevated mitochondrial H2 O2 emission during impaired oxidative phosphorylation. J Cachexia Sarcopenia Muscle 10, 643–661.

Ježek J, Engstová H & Ježek P (2017). Antioxidant mechanism of mitochondria-targeted plastoquinone SkQ1 is suppressed in aglycemic HepG2 cells dependent on oxidative phosphorylation. Biochimica et Biophysica Acta (BBA) - Bioenergetics 1858, 750–762.

Kamiya M, Mizoguchi F, Kawahata K, Wang D, Nishibori M, Day J, Louis C, Wicks IP, Kohsaka H & Yasuda S (2022). Targeting necroptosis in muscle fibers ameliorates inflammatory myopathies. Nat Commun; DOI: 10.1038/s41467-021-27875-4.

Kearney CJ, Cullen SP, Tynan GA, Henry CM, Clancy D, Lavelle EC & Martin SJ (2015). Necroptosis suppresses inflammation via termination of TNF- or LPS-induced cytokine and chemokine production. Cell Death & Differentiation 2015 22:8 22, 1313–1327.

Khoury MK, Gupta K, Franco SR & Liu B (2020). Necroptosis in the Pathophysiology of Disease. American Journal of Pathology 190, 272–285.

Kim TY, Kang JH, Lee S Bin, Kang TB & Lee KH (2021). Down-regulation of pro-necroptotic molecules blunts necroptosis during myogenesis. Biochem Biophys Res Commun 557, 33– 39.

Martin A, Gallot YS & Freyssenet D (2023). Molecular mechanisms of cancer cachexia-related loss of skeletal muscle mass: data analysis from preclinical and clinical studies. J Cachexia Sarcopenia Muscle 14, 1150.

Martin L, Birdsell L, MacDonald N, Reiman T, Clandinin MT, McCargar LJ, Murphy R, Ghosh S, Sawyer MB & Baracos VE (2013). Cancer cachexia in the age of obesity: Skeletal muscle depletion is a powerful prognostic factor, independent of body mass index. Journal of Clinical Oncology 31, 1539–1547.

Matuszewska K, Santry LA, van Vloten JP, AuYeung AWK, Major PP, Lawler J, Wootton SK, Bridle BW & Petrik J (2019). Combining vascular normalization with an oncolytic virus enhances immunotherapy in a preclinical model of advanced-stage ovarian cancer. Clinical Cancer Research 25, 1624–1638.

Morgan JE, Prola A, Mariot V, Pini V, Meng J, Hourde C, Dumonceaux J, Conti F, Relaix F, Authier FJ, Tiret L, Muntoni F & Bencze M (2018). Necroptosis mediates myofibre death in dystrophin-deficient mice. Nature Communications 2018 9:1 9, 1–10.

Mueller TC, Bachmann J, Prokopchuk O, Friess H & Martignoni ME (2016). Molecular pathways leading to loss of skeletal muscle mass in cancer cachexia – can findings from animal models be translated to humans? BMC Cancer 2016 16:1 16, 1–14.

Newton K, Sun X & Dixit VM (2004). Kinase RIP3 Is Dispensable for Normal NF-κBs, Signaling by the B-Cell and T-Cell Receptors, Tumor Necrosis Factor Receptor 1, and Toll-Like Receptors 2 and 4. Mol Cell Biol 24, 1464.

Ogilvie LM, Delfinis LJ, Coyle-Asbil B, Vudatha V, Alshamali R, Garlisi B, Pereira M, Matuszewska K, Garibotti MC, Gandhi S, Brunt KR, Wood GA, Trevino JG, Perry CGR, Petrik J & Simpson JA (2024). Cardiac Atrophy, Dysfunction, and Metabolic Impairments: A Cancer-Induced Cardiomyopathy Phenotype. Am J Pathol 194, 1823–1843.

Oikonomou N, Schuijs MJ, Chatzigiagkos A, Androulidaki A, Aidinis V, Hammad H, Lambrecht BN & Pasparakis M (2021). Airway epithelial cell necroptosis contributes to asthma exacerbation in a mouse model of house dust mite-induced allergic inflammation. Mucosal Immunology 2021 14:5 14, 1160–1171.

Penna F, Busquets S & Argilés JM (2016). Experimental cancer cachexia: Evolving strategies for getting closer to the human scenario. Semin Cell Dev Biol 54, 20–27.

Pin F, Huot JR & Bonetto A (2022). The Mitochondria-Targeting Agent MitoQ Improves Muscle Atrophy, Weakness and Oxidative Metabolism in C26 Tumor-Bearing Mice. Front Cell Dev Biol 10, 861622.

Poisson J et al. (2021). Prevalence and prognostic impact of cachexia among older patients with cancer: a nationwide cross-sectional survey (NutriAgeCancer). J Cachexia Sarcopenia Muscle 12, 1477.

Polykratis A, Hermance N, Zelic M, Roderick J, Kim C, Van T-M, Lee TH, Chan FKM, Pasparakis M & Kelliher MA (2014). RIPK1 kinase inactive mice are viable and protected from TNF-induced necroptosis in vivo. J Immunol 193, 1539.

Prado CMM, Baracos VE, McCargar LJ, Reiman T, Mourtzakis M, Tonkin K, Mackey JR, Koski S, Pituskin E & Sawyer MB (2009). Sarcopenia as a determinant of chemotherapy toxicity and time to tumor progression in metastatic breast cancer patients receiving capecitabine treatment. Clinical Cancer Research 15, 2920–2926.

Rahman FA, Hian-Cheong DJ, Boonstra K, Ma A, Thoms JP, Zago AS & Quadrilatero J (2024). Augmented mitochondrial apoptotic signaling impairs C2C12 myoblast differentiation following cellular aging through sequential passaging. J Cell Physiol; DOI: 10.1002/JCP.31155.

Rodriguez DA, Weinlich R, Brown S, Guy C, Fitzgerald P, Dillon CP, Oberst A, Quarato G, Low J, Cripps JG, Chen T & Green DR (2016). Characterization of RIPK3-mediated phosphorylation of the activation loop of MLKL during necroptosis. Cell Death Differ 23, 76.

Russell S, Duquette M, Liu J, Drapkin R, Lawler J & Petrik J (2014). Combined therapy with thrombospondin-1 type I repeats (3TSR) and chemotherapy induces regression and significantly improves survival in a preclinical model of advanced stage epithelial ovarian cancer. The FASEB Journal 29, 576.

Samson AL et al. (2020). MLKL trafficking and accumulation at the plasma membrane control the kinetics and threshold for necroptosis. Nat Commun; DOI: 10.1038/S41467-020-16887-1.

Sever AIM, Reid Alderson T, Rennella E, Aramini JM, Liu ZH, Harkness RW & Kay LE (2023). Activation of caspase-9 on the apoptosome as studied by methyl-TROSY NMR. Proc Natl Acad Sci U S A 120, e2310944120.

Skinner SK, Solania A, Wolan DW, Cohen MS, Ryan TE & Hepple RT (2021). Mitochondrial Permeability Transition Causes Mitochondrial Reactive Oxygen Species- and Caspase 3-Dependent Atrophy of Single Adult Mouse Skeletal Muscle Fibers. Cells; DOI: 10.3390/CELLS10102586.

Skulachev VP, Vyssokikh MY, Chernyak B V., Averina OA, Andreev-Andrievskiy AA, Zinovkin RA, Lyamzaev KG, Marey M V., Egorov M V., Frolova OJ, Zorov DB, Skulachev M V. & Sadovnichii VA (2023). Mitochondrion-targeted antioxidant SkQ1 prevents rapid animal death caused by highly diverse shocks. Scientific Reports 2023 13:1 13, 1–15.

Smuder AJ, Roberts BM, Wiggs MP, Kwon OS, Yoo JK, Christou DD, Fuller DD, Szeto HH & Judge AR (2020). Pharmacological targeting of mitochondrial function and reactive oxygen species production prevents colon 26 cancer-induced cardiorespiratory muscle weakness. Oncotarget 11, 3502.

Tomasin R, Martin ACBM & Cominetti MR (2019). Metastasis and cachexia: alongside in clinics, but not so in animal models. J Cachexia Sarcopenia Muscle 10, 1183.

Tsitkanou S, Morena da Silva F, Cabrera AR, Schrems ER, Muhyudin R, Koopmans PJ, Khadgi S, Lim S, Delfinis LJ, Washington TA, Murach KA, Perry CGR & Greene NP (2024). Mitochondrial antioxidant SkQ1 attenuates C26 cancer-induced muscle wasting in males and improves muscle contractility in female tumor-bearing mice. Am J Physiol Cell Physiol; DOI: 10.1152/AJPCELL.00497.2024.

Van Vledder MG, Levolger S, Ayez N, Verhoef C, Tran TCK & Ijzermans JNM (2012). Body composition and outcome in patients undergoing resection of colorectal liver metastases. Br J Surg 99, 550–557.

Yang Z, Wang Y, Zhang Y, He X, Zhong CQ, Ni H, Chen X, Liang Y, Wu J, Zhao S, Zhou D & Han J (2018). RIP3 targets pyruvate dehydrogenase complex to increase aerobic respiration in TNF-induced necroptosis. Nature Cell Biology 2018 20:2 20, 186–197.

Yow SJ, Rosli SN, Hutchinson PE & Chen KW (2024). Differential signalling requirements for RIPK1-dependent pyroptosis in neutrophils and macrophages. Cell Death & Disease 2024 15:7 15, 1–8.

Zhang X, Dowling JP & Zhang J (2019). RIPK1 can mediate apoptosis in addition to necroptosis during embryonic development. Cell Death & Disease 2019 10:3 10, 1–11.

Zhang Y, Su SS, Zhao S, Yang Z, Zhong CQ, Chen X, Cai Q, Yang ZH, Huang D, Wu R & Han J (2017). RIP1 autophosphorylation is promoted by mitochondrial ROS and is essential for RIP3 recruitment into necrosome. Nat Commun; DOI: 10.1038/ncomms14329.

