## Supplemental Figure 2 for "The mitochondrial-targeted antioxidant SkQ1 prevents mitochondrial-linked apoptosis but not necroptosis or skeletal muscle atrophy in ovarian cancer"

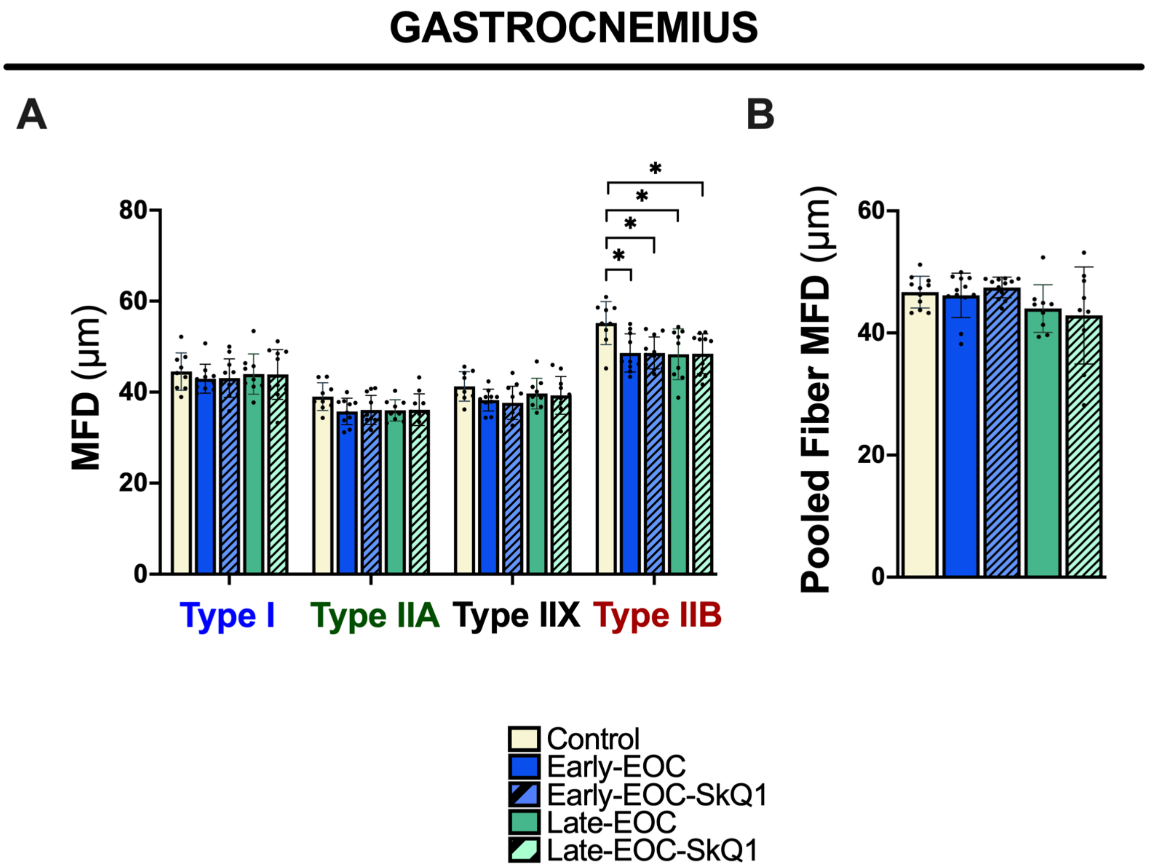


**Supplemental Figure 2. Evaluation of gastrocnemius fiber type atrophy with ovarian cancer progression. A)** Minimal-feret diameter (MFD) of gastrocnemius muscle fibers stained for myosin-heavy chain (MHC) isoforms and **B)** pooled MFD of gastrocnemius fibers, n=9 -12. Analyses for **A and B** were conducted using either a standard one-way ANOVA or Kruskal-Wallis test, depending on whether data was normally distributed. These were followed by a two-step step-up method of Benjamini, Krieger, and Yekutieli for post-hoc analysis. Results represent mean ± SD. ***** = $p < 0.05$.
