## Supplemental Figure 3 for "The mitochondrial-targeted antioxidant SkQ1 prevents mitochondrial-linked apoptosis but not necroptosis or skeletal muscle atrophy in ovarian cancer"

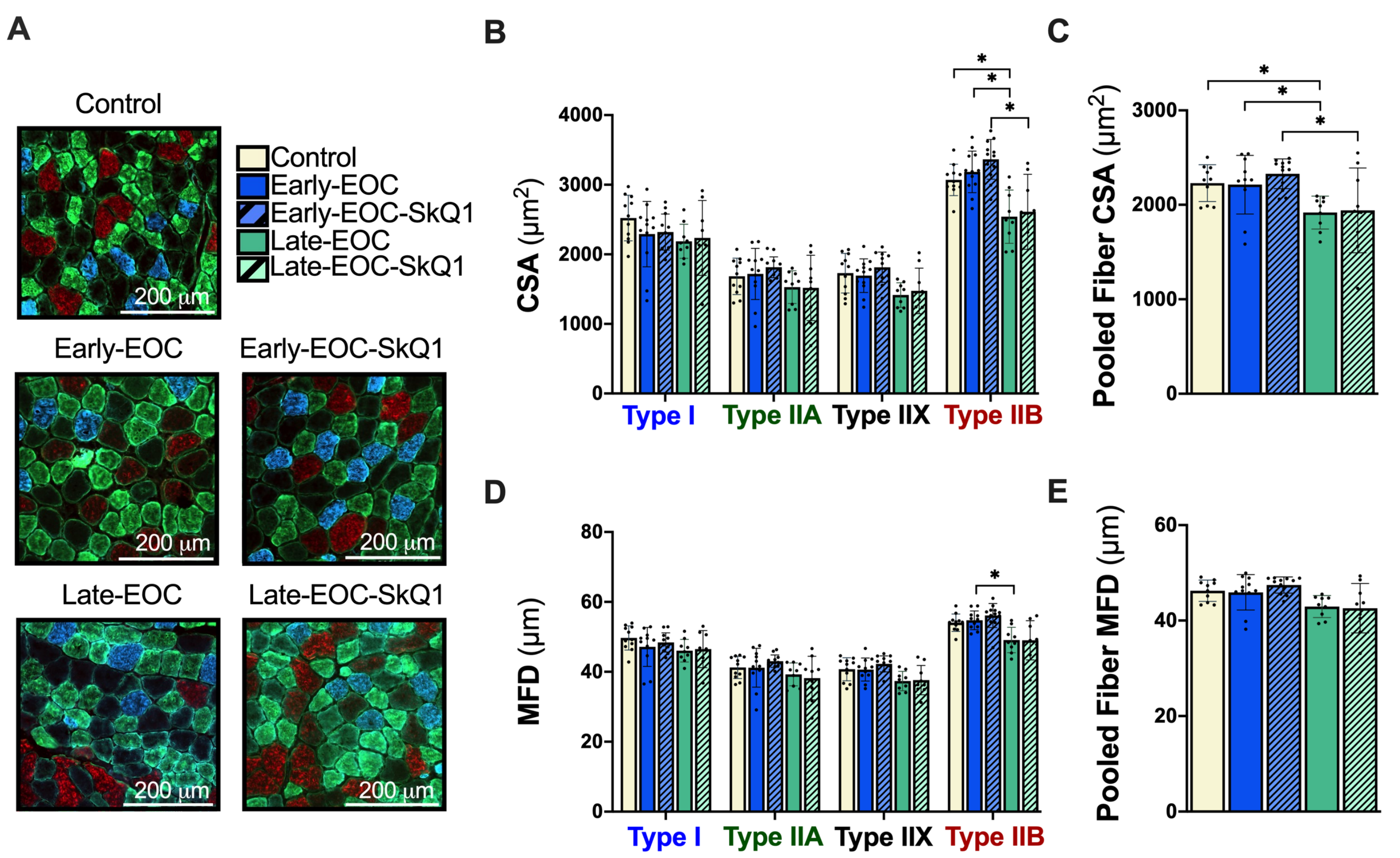


**Supplemental Figure 3. Quadriceps atrophy with EOC progression. A)** Representative images of quadriceps cross-sections stained for myosin heavy chain (MHC) isoforms. Blue corresponds to type I fibers, green: type IIA, black: type IIX, and red: type IIB fibers. **B)** Cross-sectional area (CSA) of quadriceps muscle fibers (n=9 -12). **C)** Pooled quadriceps muscle fiber CSA. **D)** Minimal-feret diameter (MFD) of quadriceps muscle fibers and **E)** pooled MFD of quadriceps fibers (n = 9-12). Analyses for **B**, **C**, **D**, and **E** were conducted using either a standard one-way ANOVA or Kruskal-Wallis test, depending on whether data was normally distributed. These were followed by a two-step step-up method of Benjamini, Krieger, and Yekutieli for post-hoc analysis. Results represent mean ± SD. * = $p < 0.05$.
