## Supplemental Figure 4 for "The mitochondrial-targeted antioxidant SkQ1 prevents mitochondrial-linked apoptosis but not necroptosis or skeletal muscle atrophy in ovarian cancer"

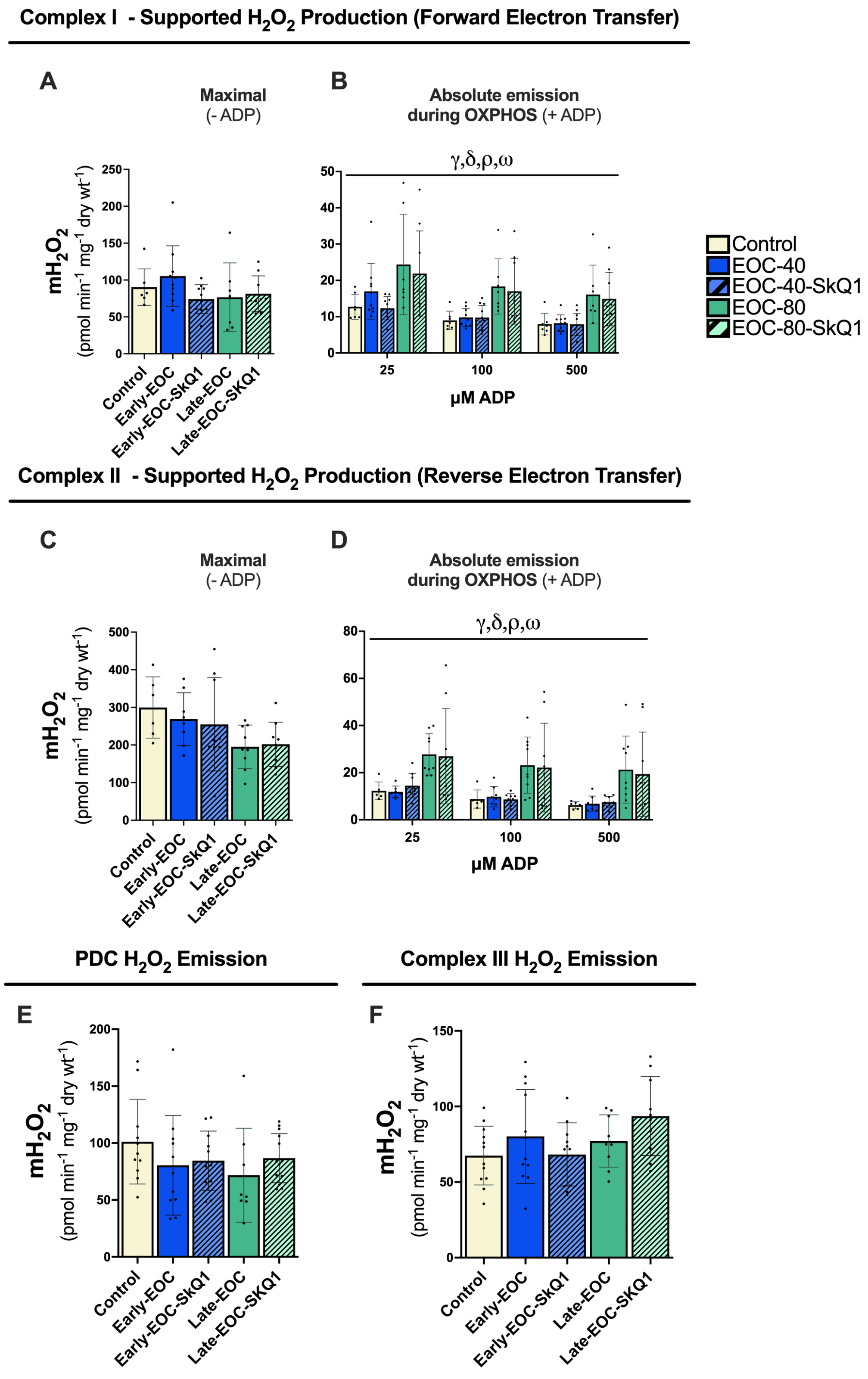


**Supplemental Figure 4. Site-specific $mH_2O_2$ emission in the white gastrocnemius of EOC-tumour bearing mice subjected to either SkQ1 or normal drinking water. A)** Complex I-supported (NADH supplied via pyruvate and malate) maximal $mH_2O_2$ (in the absence of ADP; -ADP (n= 7-11). **B)** Absolute rates of Complex I - supported $mH_2O_2$ in the presence of ADP (+ADP) to model kinetics during oxidative phosphorylation. (n=7-11). **C)** Complex II-supported ($FADH_2$ supplied via succinate) maximal $mH_2O_2$ (-ADP). (n=7-11) **D)** Absolute rates of Complex II-supported $mH_2O_2$ in the presence of ADP (+ADP) to model kinetics during oxidative phosphorylation. **E)** Pyruvate dehydrogenase complex-derived (pyruvate and rotenone) maximal $mH_2O_2$ (n= 8-13). **F)** Complex III derived (Antimycin A) maximal $mH_2O_2$ (n = 10-12). **$H_2O_2$**: Hydrogen Peroxide. Analyses for **A**, **C**, **E** and **F** were conducted using either a standard one-way ANOVA or Kruskal-Wallis test, depending on whether data was normally distributed. Analyses for **B** and **D** were conducted using a standard two-way ANOVA. Analyses were followed by a two-step step-up method of Benjamini, Krieger, and Yekutieli for post-hoc analysis. Results represent mean ± SD. ***** = $p < 0.05$. **γ** = $p < 0.05$ Control vs Late-EOC, **δ** = < 0.05 Control vs Late-EOC-SkQ1, **ρ** = $p < 0.05$ Early-EOC vs Late-EOC, **ω** = Early-EOC-SkQ1 vs Late-EOC-SkQ1.
