## Supplemental Figure 5 for "The mitochondrial-targeted antioxidant SkQ1 prevents mitochondrial-linked apoptosis but not necroptosis or skeletal muscle atrophy in ovarian cancer"

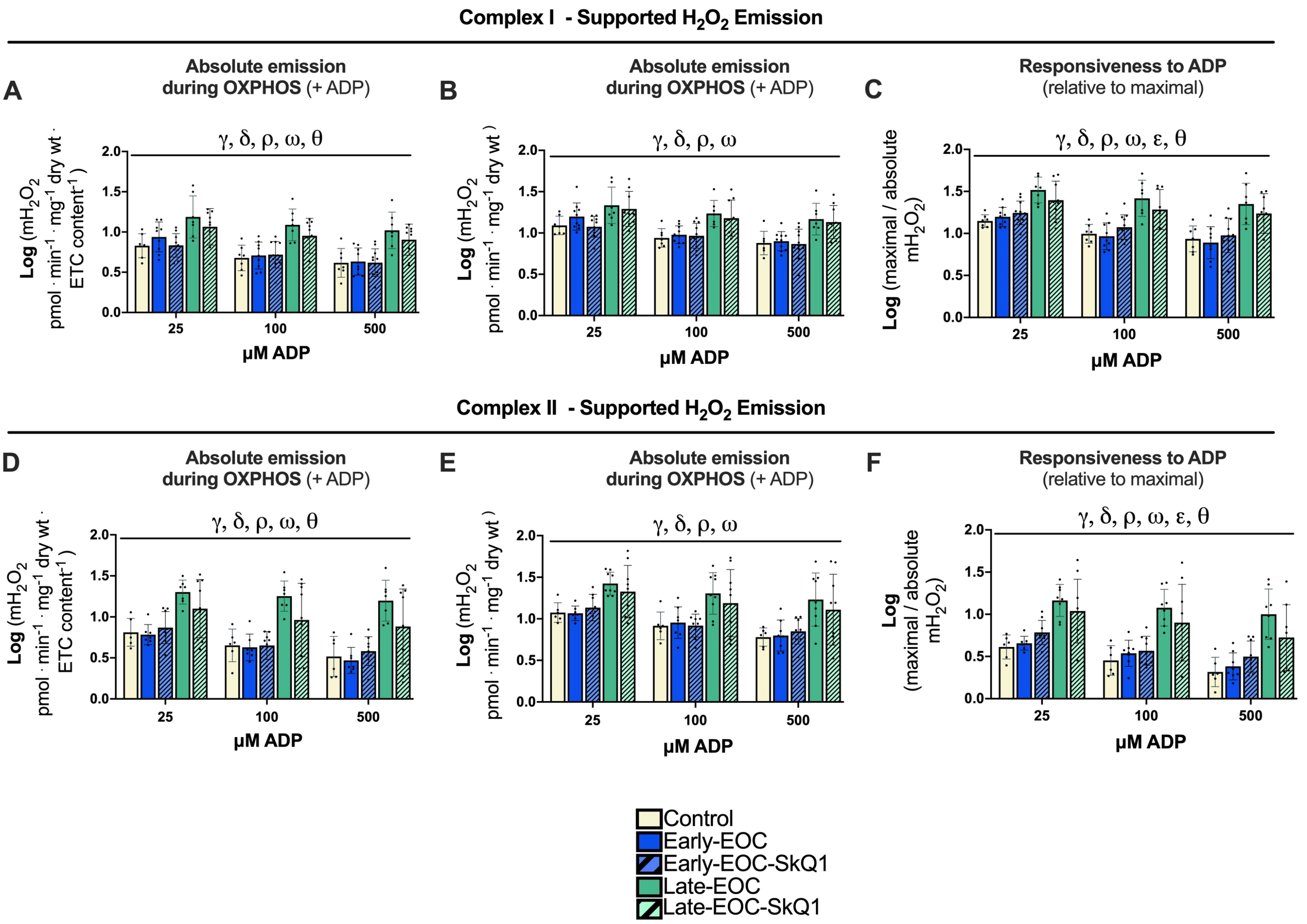


**Supplemental Figure 5. Log transformed $H_2O_2$ emission data**. **A)** Absolute rates of Complex I -supported $mH_2O_2$ in the presence of ADP (+ADP) normalized to ETC content (n=7-11). **B)** Absolute rates of Complex I -supported $mH_2O_2$ in the presence of ADP (+ADP, n=7-11). **C)** Complex-I $mH_2O_2$ during oxidative phosphorylation (+ADP) expressed relative to maximal (-ADP) rates as an index of mitochondrial responsiveness to ADP's attenuating effects (n=7-11). **D)** Absolute rates of Complex II-supported $mH_2O_2$ in the presence of ADP (+ADP) to model kinetics during oxidative phosphorylation normalized to ETC content (n=7-11). **E)** Absolute rates of Complex II-supported $mH_2O_2$ in the presence of ADP (+ADP) to model kinetics during oxidative phosphorylation. **F)** Complex-II $mH_2O_2$ during oxidative phosphorylation (+ADP) expressed relative to maximal (-ADP) rates as an index of mitochondrial responsiveness to ADP's attenuating effects (n=7-11). All data were non-normally distributed and were first log-transformed, then analyzed using a standard two-way ANOVA. Analyses were followed by a two-step step-up method of Benjamini, Krieger, and Yekutieli for post-hoc analysis. Results represent mean ± SD. ***** = $p < 0.05$. **γ** = $p < 0.05$ Control vs Late-EOC, **δ** = < 0.05 Control vs Late-EOC-SkQ1, **ε** = $p < 0.05$ Early-EOC vs Early-EOC-SkQ1, **θ** = $p < 0.05$ Late-EOC vs Late-EOC-SkQ1, **ρ** = $p < 0.05$ Early-EOC vs Late-EOC, **ω** = Early-EOC-SkQ1 vs Late-EOC-SkQ1.
