## Supplemental Figure 6 for "The mitochondrial-targeted antioxidant SkQ1 prevents mitochondrial-linked apoptosis but not necroptosis or skeletal muscle atrophy in ovarian cancer"

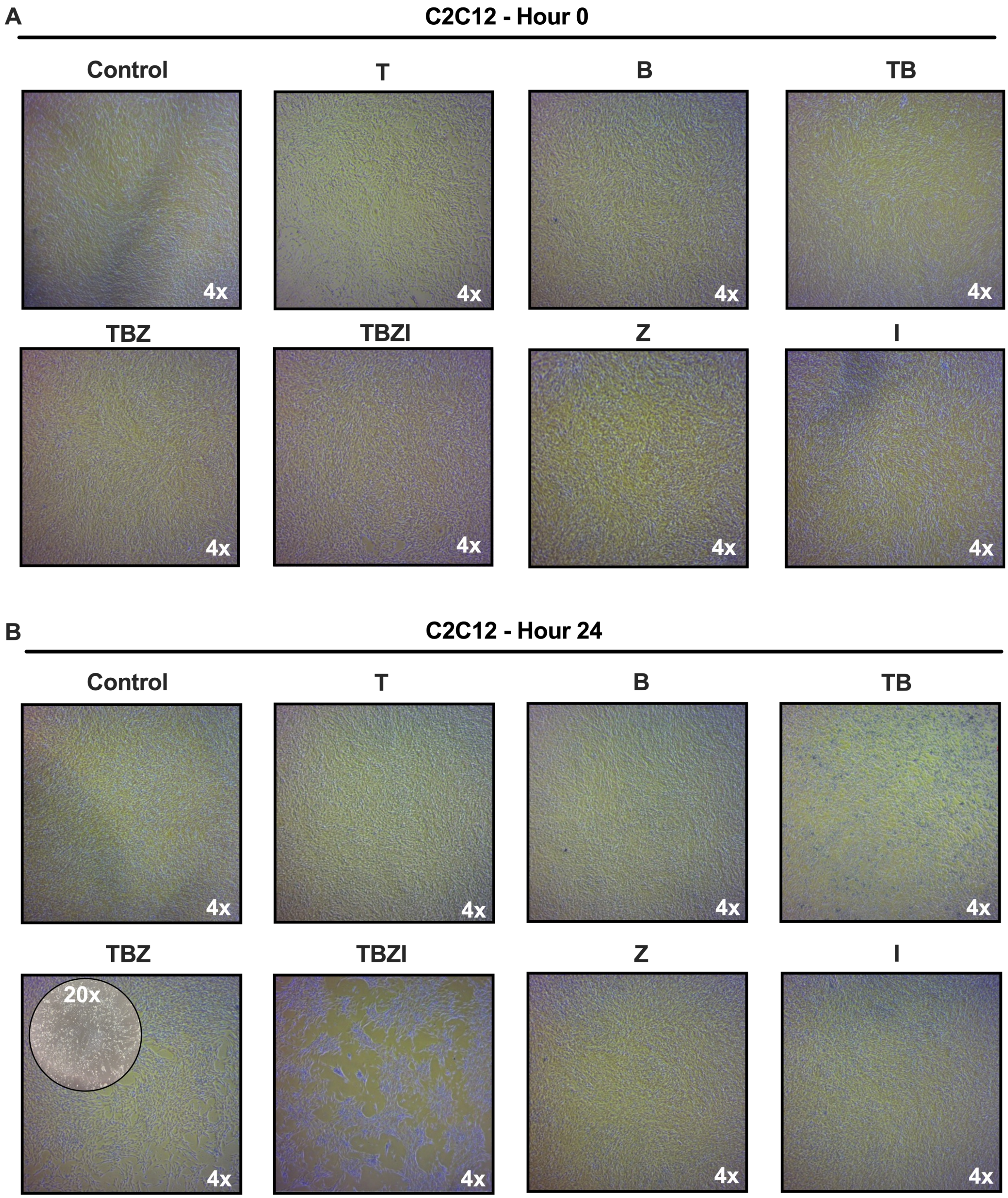


**Supplemental Figure 6. C2C12 myoblasts pre (hour 0) and post (hour 24) combination of TNF-α (T), B (BV6), z-VAD-FMK (Z) and z-IEDT-FMK (I) treatment**. **A)** Images of C2C12 plates captured at 4x magnification prior to TBZI treatment. **B)** Images of C2C12 plates captured at 4x magnification 24 hours post TBZI treatment. BSA (T vehicle) and DMSO (BZI Vehicle) plates not shown.
