## Supplemental Figure 8 for "The mitochondrial-targeted antioxidant SkQ1 prevents mitochondrial-linked apoptosis but not necroptosis or skeletal muscle atrophy in ovarian cancer"

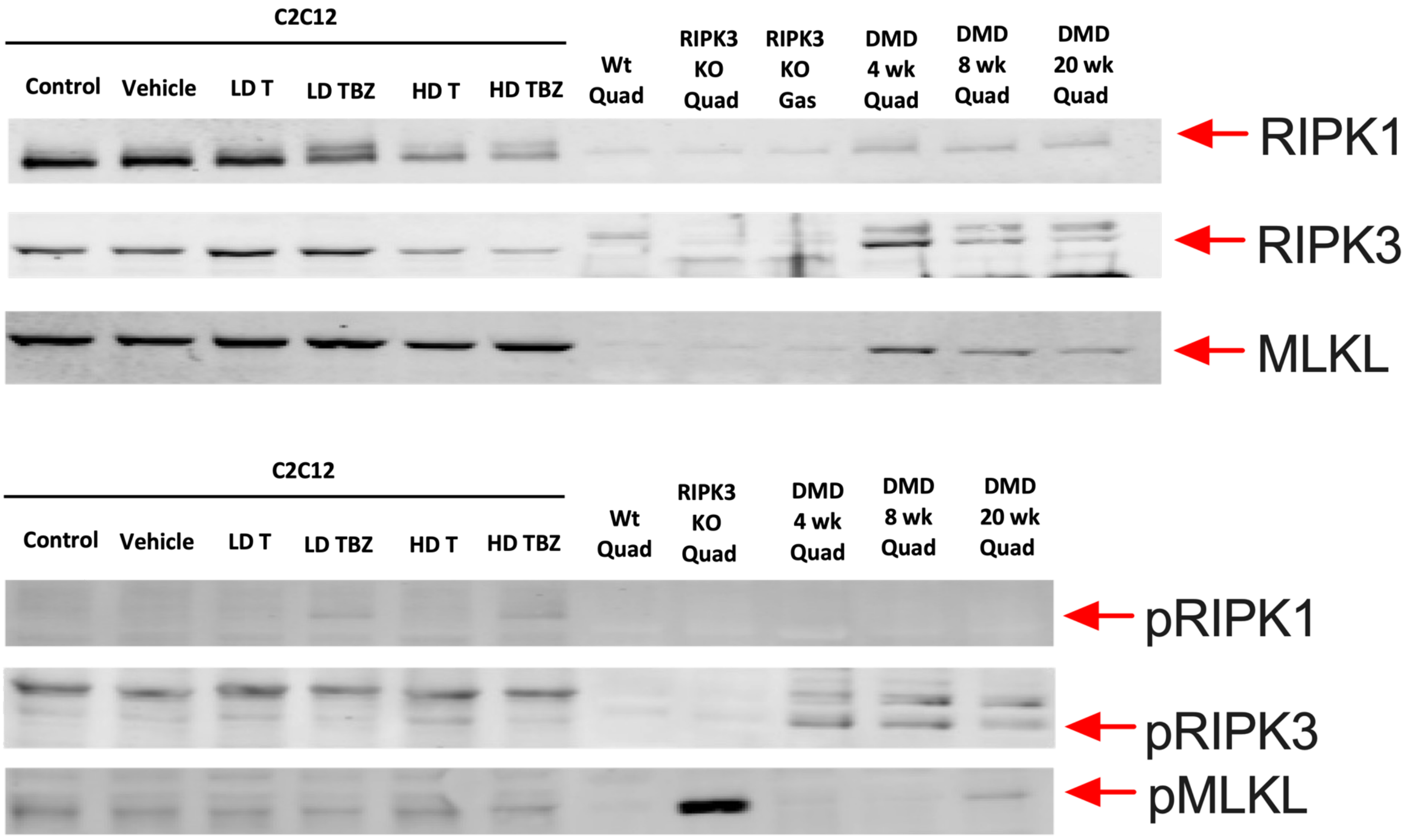


**Supplemental Figure 8. Western blots of C2C12s treated with different combinations of TBZI, wild-type quadriceps, RIPK3 KO quadriceps and gastrocnemius, and quadriceps from D2.mdx mice at different ages. Vehicle:** BSA (TNF-α vehicle) and DMSO (BV6 and z-Vad-FMK vehicle). **LD T:** low dose TNF-α. **LD TBZ**: low dose TNF-α, BV6 and z-Vad-FMK. **HD T:** high dose TNF-α. **HD TBZ**: high dose TNF-a, BV6 and z-Vad-FMK. **Wt**: wild-type. **RIPK3 KO**: RIPK3 knockout. **DMD**: Duchenne muscular dystrophy. **Quad**: quadriceps. **Gas**: gastrocnemius. **RIPK1**: Receptor Interacting Serine/Threonine Protein Kinase 1, **RIPK3**: Receptor Interacting Serine/Threonine Protein Kinase 3, **MLKL**: Mixed-Lineage Kinase Domain-Like Protein, **pRIPK1**: phosphorylated RIPK1, **pRIPK3**: phosphorylated RIPK3, **pMLKL**: phosphorylated MLKL.
