## Supplemental Figure 7 for "The mitochondrial-targeted antioxidant SkQ1 prevents mitochondrial-linked apoptosis but not necroptosis or skeletal muscle atrophy in ovarian cancer"

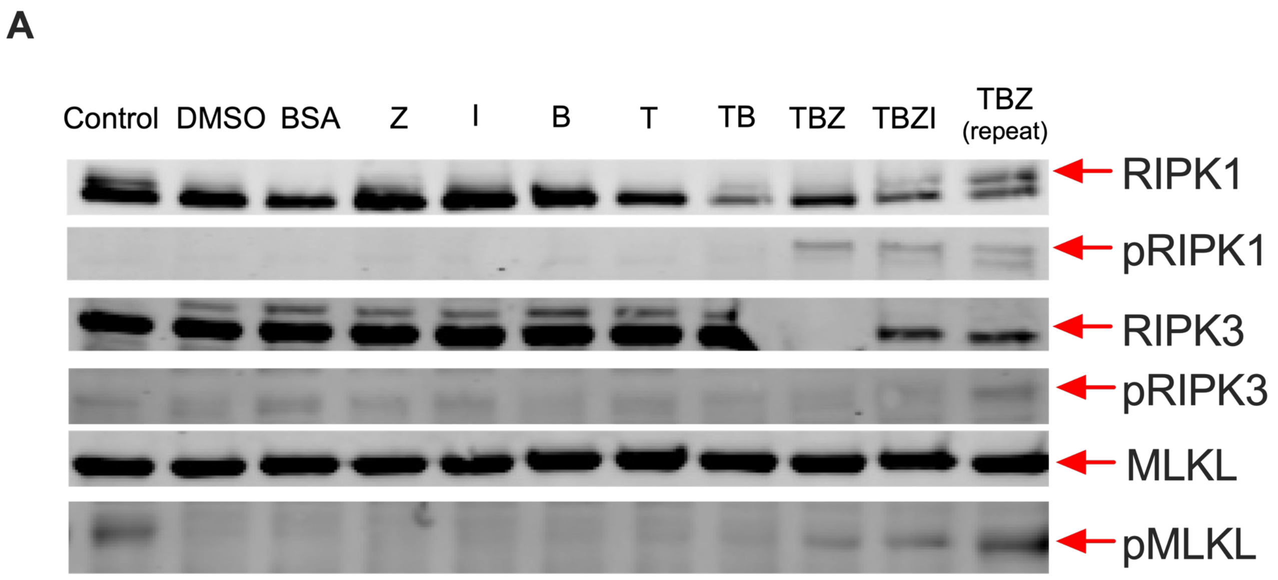


**Supplemental Figure 7**. **Western blots of C2C12s treated with different combinations of TBZI. RIPK1**: Receptor Interacting Serine/Threonine Protein Kinase 1, **RIPK3**: Receptor Interacting Serine/Threonine Protein Kinase 3, **MLKL**: Mixed-Lineage Kinase Domain-Like Protein, **pRIPK1**: phosphorylated RIPK1, **pRIPK3**: phosphorylated RIPK3, **pMLKL**: phosphorylated MLKL.
