## Supplemental Figure 1 for "The mitochondrial-targeted antioxidant SkQ1 prevents mitochondrial-linked apoptosis but not necroptosis or skeletal muscle atrophy in ovarian cancer"

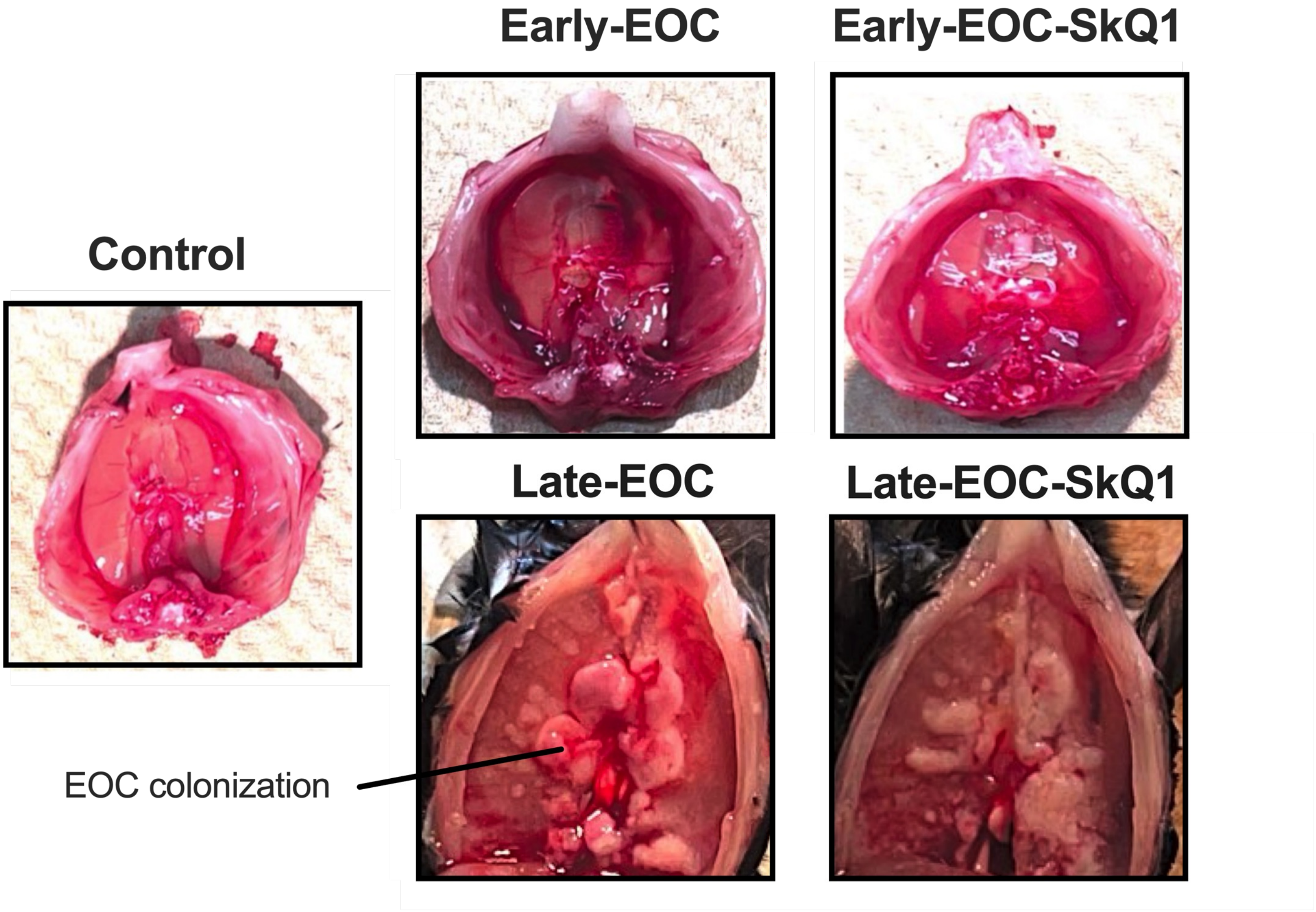


**Supplemental Figure 1. Inferior view of diaphragm muscle from ovarian tumour bearing mice.** White areas on the diaphragm are populations of invading EOC cells. Photograph for Early-EOC has been previously published (Delfinis *et al*, 2024) and re-used with permission by the authors of this manuscript in accordance with Elsevier policy on copyright pertaining to 'author rights'.
